# habtools: an R package to calculate 3D metrics for surfaces and objects

**DOI:** 10.1101/2024.09.19.613985

**Authors:** Nina Schiettekatte, Mollie Asbury, Guan-Yan Chen, Maria Dornelas, Jessica Reichert, Damaris Torres-Pulliza, Kyle J.A. Zawada, Joshua S. Madin

## Abstract

1. Technological advances in three-dimensional imaging techniques have opened the door to advanced morphological analyses and habitat mapping for biologists and ecologists.
2. At the same time, the challenge of translating complex 3D data into meaningful metrics that can be used in conjunction with biological data currently hinders progress and accessibility.
3. We introduce *habtools*, an R package that provides R functions to efficiently calculate complexity and shape metrics from DEMs, 3D meshes and 2D shapes as well as some helper functions to facilitate workflow.
4. We expect the functionality of *habtools* to continue to expand as new metrics and faster methods become available, and welcome new contributions and ideas.

## 1. Introduction

Structural complexity refers to how the surface and volume of an object or a habitat is distributed in space, and can explain phenomena on the individual organism level and biodiversity and ecosystem functioning patterns on the community level (LaRue *et al*. 2023). Habitat complexity has been shown to augment abundance and species richness (Graham & Nash 2013; Torres-Pulliza *et al*. 2020), and influence size distributions of animals inhabiting the structure, which may indirectly affect energy fluxes within the ecosystem (Nash *et al*. 2013). For example, the fractal dimension of habitat affects the body sizes of arthropods living on plants (Morse *et al*. 1985; Gunnarsson 1992) and fishes living on coral reefs (Nash *et al*. 2013). Moreover, the shape of habitat elements can also affect their vulnerability to storms (Madin & Connolly 2006). Whilst links between shape and ecology have been demonstrated, effectively assessing the role of highly complex structures on ecological patterns and processes hinges upon our ability to quantify it.

Recent technological advances such as photogrammetry, structure-from-motion (SfM) techniques, light detection and ranging (LiDAR), and 3D scanning can generate fine-detailed 3D reconstructions of individual organisms and complex habitats alike, such as coral reefs or forests (Ferrari *et al*. 2016; Dubayah *et al*. 2020). These techniques result in the generation of different datasets representing a surface or volume: 3D point clouds are collections of data points in three-dimensional space; 3D meshes are collections of interconnected polygons, typically triangles and consisting of vertices, edges, and polygonal faces; digital elevation models (DEMs) are collections of height information in the form of a raster grid or a triangular irregular network. Based on these datasets, simple 3D metrics, like surface area, rugosity, height, or the number and size of holes and crevices can be derived (Fukunaga & Burns 2020; Seidel *et al*. 2021). Beyond this, different methods to describe the form of habitats have been developed, such as fractal dimension, top-heaviness, or convexity (Reichert *et al*. 2017; Zawada *et al*. 2019; Million *et al*. 2021; Aston *et al*. 2022). While some metrics are straightforward in their estimation (e.g. rugosity), metrics such as fractal dimension are harder to estimate due to methodological challenges, potential sources of bias, and computation time. Given the importance of shape across systems and scales, and the proliferation of both methods and data to quantify it, there is a need for accessible and standardized software to facilitate data integration, interpretation, and analysis.

Different approaches to quantify structural metrics have been compiled in software packages or published code. For example, algorithms for the analysis of 3D meshes of corals have been implemented in python/meshlab (Fukunaga *et al*. 2019; Aston *et al*. 2022), and point clouds and quantitative structure models of trees can be analyzed using the R packages `FORTLS` (Molina-Valero *et al*. 2022) or `ITSMe` (Terryn *et al*. 2023). Furthermore, the python package `HSC3D` provides specialized algorithms to calculate complexity metrics for intertidal habitats based on photogrammetry point clouds (Gu *et al*. 2024). While specialized snippets of code and organism or habitat-specific packages have been developed to aid the analysis of 3D data, there is currently no software package that aids with the analysis of different types of 3D data in the R environment.

Here, we present the R package `habtools` that includes a collection of R functions to efficiently calculate complexity and shape metrics from DEMs, 3D meshes and 2D shapes as well as some helper functions to facilitate workflow and diagnose fractal dimension calculations. We introduce a workflow using `habtools` and provide an in-depth discussion on the calculation of fractal dimension, a metric that requires a conscious user-defined choice of method and scale. We further present two case studies centered around corals and coral reefs, however, the methods can be applied to any 3D object or surface.

## 2. Description habtools

The goal of `habtools` is to provide a set of user-friendly functions to calculate 3D metrics for a range of surfaces, i.e. digital elevation models (DEM) and 3D meshes of objects, targeted at biologists and ecologists. The metrics can be used for any object or surface, ranging from single cells (e.g. microglia brain cells; Salamanca *et al*. (2019)) to large habitat patches (e.g. forest canopy; Hardiman *et al*. (2013))(See Table 1 for examples). While some functions (i.e. surface area, rugosity) are applicable to both 3D objects and surfaces, a set of functions related to object shape is only available to use for 3D objects (Figure 1, Table 1). Aside from functions to calculate metrics, we also include functions to aid processing of DEMs, as illustrated in two case studies. Lastly, we provide introduction vignettes demonstrating `habtools` functions for DEMs and meshes.

**Table 1.** Description and application examples of the complexity and shape metrics, currently included in habtools.

| Metric | Units and range | Description | Example |
| --- | --- | --- | --- |
| Convexity | Unitless, ranging between 0 and 1. | Measures the ratio of the volume of an object to the volume of its convex hull. A value closer to 1 indicates a nearly convex shape, while lower values indicate more concave shapes. | Used to classify microglia cells in injured brains (Salamanca *et al.* 2019). |
| Packing | Unitless ratio centered around 1. | Ratio of the surface area of an object to the surface area of its convex hull, reflecting how much an object fills its convex hull. Values of 1 indicate low surface complexity, values above 1 indicate complex surfaces within the bulk of the volume, and values below 1 indicate complex surfaces outside of the bulk of the volume. | Coral assemblage-scale morphological traits, including packing, have been shown to be affected by disturbance events in coral reefs (Zawada *et al.* 2019). |
| Sphericity | Unitless, ranging between 0 and 1. | Quantifies how closely an object's shape resembles a sphere. Values closer to 1 indicate more spherical and compact shapes. | Affects pollen grain aerodynamics and wind dispersal in plants (Jackson & Lyford 1999). |
| Top-heaviness (SMA, SMV) | SMA: length unit<sup>3</sup> from 0 (e.g. cm<sup>3</sup>); SMV: length unit<sup>4</sup> from 0 (e.g. cm<sup>4</sup>) | Measures an object's distribution of area (SMA) or volume (SMV) relative to an axis, providing insight into how "top-heavy" a structure is. | Top-heaviness of tree crowns predict respiration rates (Röll *et al.* 2023). |
| Mechanical Shape Factor (CSF) | Positive, dimensionless value. | Quantifies the mechanical vulnerability of a structural element, describing how an object's shape influences its ability to resist cantilever drag forces. | Can be used to predict the effects of hydrodynamic disturbances on coral reef communities (Madin & Connolly 2006). |
| Rugosity | Dimensionless value, higher than or equal to 1. | A measure of surface roughness or texture, often used to quantify the structural complexity of surfaces. Calculated as the ratio of surface area to planar area. | Canopy rugosity can be used as a robust proxy of stand carbon storage potential in forests (Hardiman *et al.* 2013). |
| Fractal Dimension | Dimensionless value, typically ranging between 2 and 3 for 3D shapes and 1 and 2 for 2D shapes. | Estimates the space filling capacity of a surface or line. A higher fractal dimension indicates more complex structures. | The fractal dimension of plants affects the body sizes of arthropods living on them (Morse *et al.* 1985). |
| Height Range | Positive length unit (e.g. cm). | Measures the vertical distribution of elements within a structure. Calculated as the maximum height subtracted by the minimum height. | Can be used to determine tree heights from 3D data obtained using airborne lidar (Andersen *et al.* 2006). |

**Figure 1.**
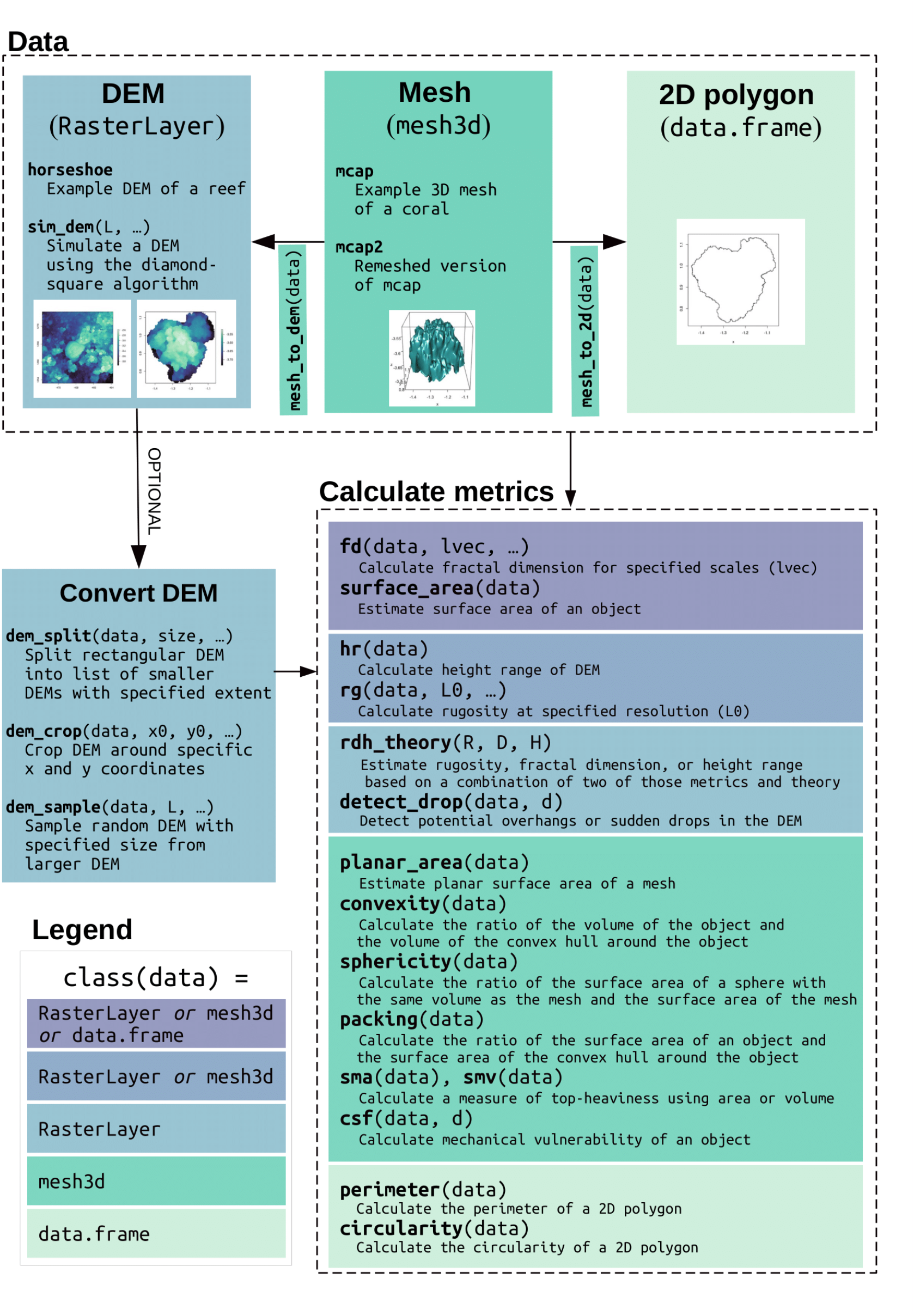
Overview of main habtools functions and workflow. Functions are grouped by the input data type.

### 2.1 Workflow

How long is a piece of string? Before starting any metric calculations, it is crucial to consider if the resolution of the objects or surfaces at hand matches the scale of the phenomena of interest. For example, if we want to test the effect of landscape rugosity on the density of elephants, we may want to avoid a mm-scale resolution that is not relevant for the elephant’s ecology and instead, use a coarser resolution for this question.. The resolution of choice will impact calculations and may yield varying results. We advise ensuring that (1) all objects or surfaces have the same resolutions when comparing metrics across objects, and (2) the resolution suits the scientific question. Higher resolution is not always better. Once the resolution is carefully chosen and homogenized across objects, `habtools` can be used to transform 3D data and calculate complexity and shape metrics (Figure 1, Table 1)

For DEMs, users can load and transform a RasterLayer using `habtools` functions. The RasterLayer can be used as is, however, `dem_crop()` allows users to cut out a square-shaped DEM around a specific point of interest and of a certain size. If within-site variation of complexity metrics is of interest, one can use the `dem_split()` function to subdivide a large square-shaped DEM into smaller square-shaped DEMs of a desired size. One can also crop out a randomly placed DEM of a given size using `dem_sample()`, which has the option to exclude data gaps (i.e. where z data is missing). Once a square DEM or list of DEMs of desired size is obtained, complexity metrics can be calculated. There are currently three main metrics included for DEMs (Torres-Pulliza *et al*. 2020): height range (i.e. the distance between highest and lowest points, `hr()`), rugosity (i.e. the ratio between surface area and planar area, `rg()`), and fractal dimension (i.e. a metric describing how a surface fills a volume, `fd()`). Finally, the function `detect_drop()` allows users to identify sudden height changes or potential overhangs in a DEM, which can be useful for diagnosing 3D metrics or indicate potential hiding spaces for mobile animals.

For working with meshes, users should ensure a rectified orientation (i.e. z axis is aligned with the up facing direction) and equal resolution of the mesh. If the resolution (i.e. spacing between vertices) varies throughout the mesh, it is advisable to remesh with a fixed resolution as preparation step, either in R (e.g. using the R package `Rvcg`, Schlager (2017)) or by using a 3D software (e.g. meshlab or blender). Once the mesh is checked and loaded, `habtools` provides functions to calculate complexity and shape metrics (See Figure 1, Table 1). To calculate fractal dimension (`fd()`), the cube-counting method is preferred and most widely used (See 2.2). Finally, `habtools` provides a set of conversion functions, including a function to convert meshes to 2D outlines alongside functions to calculate 2D metrics, including 2D fractal dimension, circularity, and perimeter length.

### 2.3 Fractal dimension methods

Fractal dimension is a measure of complexity describing the space-filling capacity of a surface. This metric is an ecologically-relevant and has been linked to biodiversity and size distributions of mobile animals Nash *et al*. (2013). However, there are some challenges associated with calculating fractal dimension. First, most surfaces or 3D objects are not perfect fractals (i.e. they do not behave similarly across scales), and second, even if a surface is a fractal, each method to approximate the fractal dimension has drawbacks. Whether or not one should use fractal dimension despite existing challenges has been discussed elsewhere (Loke & Chisholm 2022; Madin *et al*. 2023; Fischer & Jucker 2024). Here, we focus on providing the tools to calculate and investigate fractal dimension using multiple methods and provide an overview of potential biases and recommendations.

Generally, fractal dimension is estimated by measuring values (e.g., counts of cubes, heights, areas) across a sequence of different length scales (`lvec`), and then finding the slope between the scales and the measured values. `habtools` currently provides four methods to estimate fractal dimension in 3D: the height variation method (e.g. Torres-Pulliza *et al*. 2020), the standard deviation method (e.g. Fischer & Jucker 2024), the area method (Fukunaga *et al*. 2019), and the cube-counting method (Zawada *et al*. 2019) (Table 1, Supplementary figure 1).

The height variation method (`method = “hvar”`) is based on the relationship between changes in height range and the scale of observation. The DEM is divided into squares with side lengths specified as a vector, `lvec`. For each size, the height range of each sub-square is calculated and then the mean or median of all squares is obtained This results in a data frame with scale and the average height range estimates. The fractal dimension is then calculated as: *D*=3 −*b*, with *b* being the slope of the linear regression between the log transform of height range (*h*) and scale (*l*): *l o g* (*h*)∼ *l o g* (*l*) (Supplementary figure 1 C, D). Because a range of values is needed to calculate a height range, the smallest possible value of `lvec` should always be at least two times larger than the resolution of the DEM. The largest value in `lvec` is generally equal to the extent of the DEM. We suggest choosing a vector of doubling values for `lvec`. Importantly, the method is sensitive to the resolution of the DEM (Supplementary figure 4). Therefore, it is crucial to always use raster data with the same resolution when comparing multiple surfaces.

The standard deviation method (`method = “sd”`) is similar to the height variation method. It uses the standard deviation of heights instead of height ranges. The fractal dimension is calculated as *D*=3 *– b*, with *b* being the slope of the linear regression between log-transformed standard deviations (*sd*) and log-transformed scales (*l*) (Supplementary figure 1 E,F). While the height variation method can be very sensitive to outliers in height, the standard deviation is less sensitive and may be more robust (Fischer & Jucker 2024). We suggest a similar choice of `lvec` when using the standard deviation method as with the height variation method, discussed above.

The area method (`method = “area”`) estimates surface areas at varying scales and calculates the fractal dimension as: *D*=2−*b*, where b is the slope of the linear regression between surface area (*a*) and scale (*l*): *l o g* (*a*) ∼ *lo g* (*l*) (Supplementary figure 1 I,J). When using DEMs, surface area is calculated at each scale by projecting the DEM to the resolution of that scale (using `terra::project()`, Hijmans (2024)), as opposed to aggregating (e.g. Fukunaga *et al*. 2019) to provide more flexibility in the choice of the scales that can be included. The area method is highly sensitive to the height range (Supplementary figure 4A). When the height range is very low, relative to the extent, or varies a lot across DEMs, it is recommended to rescale the heights to match the extent (e.g. by adding `scale = TRUE`) when using the area method (Fischer & Jucker 2024). The `area` method for 3D meshes uses `Rvcg` (Schlager 2017) to remesh the object at varying resolutions, which works best on meshes that are watertight and when avoiding scales that approach the extent. The `area` method for DEMs has increased bias with increasing values inside `lvec`, because surface area is estimated for each cell by using the height of the cell and its 8 surrounding cells to create a surface (using the `surfaceArea` function from the `sp` package, Jenness (2004)). Therefore, the cells outside of the DEM are assumed to be of equal height as the border cells, resulting in a lower surface area. If the ratio of border cells to total cells inside a DEM is high, there will be an underestimation of surface area, which is the case when values inside `lvec` are too close to the extent. Therefore, the `area` method may be better suited for larger areas, relative to the chosen `lvec`. We recommend a maximum scale of 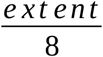 to be included inside `lvec`.

The cube-counting method (`method = “cubes”`) counts the number of cubes that intersect with a surface or outer shell of 3D object using varying cube sizes or scales. The fractal dimension is calculated as: *D*=−*b*, where *b* is the slope of the linear regression between log-transformed number of cubes that overlap with the surface (*n*) and log-transformed scales (*l*): *l o g* (*n*)∼ *l o g* (*l*) (Supplementary figure 1 G,H). This is the most commonly used method for 3D meshes and works best for closed, complex objects. When specifying `lvec`, the smallest scale should be bigger than the resolution to ensure no cubes fall in between two 3D points. While the largest scale can be equal to the extent, it may be best to exclude larger scales when the object fills the cube’s bounding box. For example, a perfect sphere with a diameter of 1 will occupy all cubes at a scale of 1 and a scale of 0.5. The fractal dimension between those large scales will then be equal to 3 ((*l o g* (1)− *lo g* (8))/*l o g* (0.5)). On those scales, a sphere fills the volume very well, while on smaller scales the fractal dimension of a sphere approximates 2. Therefore, caution should be used when including larger scales and using the cube-counting method. We further include the box-counting method (`method = “boxes”`) for 2D shapes, which is the 2D equivalent of the cube-counting method.

Deciding which method and sensible `lvec` to use can be a daunting task. To assist with this decision, `habtools` provides diagnostics to investigate a method. Specifically, the function `fd_diagnose()` renders a plot illustrating the variation across scales, and provides a list with fractal dimension across scales, the mean fractal dimension, and the standard deviation of fractal dimensions across scales (This output can also be created by setting `diagnose = TRUE` inside `fd()`). Selecting a sensible sequence of scales (i.e., the length vector `lvec`) is critical and should be kept the same when comparing surfaces or objects. Choosing a meaningful `lvec` is particularly important for non-fractal surfaces and objects as the rate of change or the fractal dimension can vary across scales. In fact, varying fractal dimensions across neighboring scales is a common phenomenon in nature (e.g. Martin-Garin *et al*. (2007)), and the diagnostic plot can help identify such transitions in rate of change. Furthermore, the best method for fractal dimension calculation may depend on the type of object and investigating the diagnostic plot may help to identify the method with lower variation. The height variation method and the standard deviation method can only be used for DEMs while the area method and the cube-counting method can be used for both DEMs and meshes. When applying various methods to the same DEMs, estimates typically correlate positively, but there can be considerable variation across methods (see supplementary materials, Supplementary figure 2). When comparing estimated fractal dimensions with theoretical values of simulated DEMs (using fractional Lévy motions), we found the standard deviation method to be most accurate (Supplementary figure 3), however, it remains to be seen if this is true for natural DEMs.

## 3. Case studies

### 3.1 Case study 1: The effect of rugosity and fractal dimension on coral diversity

Both rugosity and fractal dimension have been shown to explain variation in coral species richness, expressed as the number of unique species observed (Torres-Pulliza *et al*. 2020). Corals are ecosystem engineers so the link between structure and diversity could be related to the fact that corals provide structure. For this case study, we investigate how the relationship between complexity metrics and coral diversity varies when considering visible corals versus cryptic corals. We use the reef structure and coral diversity data examined in Torres-Pulliza *et al*. (2020). Using Structure-from-motion, a digital elevation model (DEM) of a large reef patch (∼562m^2^) near Lizard Island, Australia was constructed. All corals were annotated and identified in the field, resulting in a shape file of annotated dots that can be spatially matched with the DEM. The annotated corals included both visible corals (i.e. seen from a top-down view) and cryptic corals (i.e. hidden under a ledge or in a crevice). Here, we illustrate how `habtools` can be used to link structural complexity and biodiversity (see supplementary materials for a detailed description of the methods and code). Specifically, we compare how structural complexity metrics (rugosity and fractal dimension) correlate with visible corals and cryptic corals, respectively.

We use the `dem_sample()` function to sample 500 square DEMs with an side of 1m out of the larger reef patch. For each smaller square (1m^2^), we quantified rugosity and fractal dimension using functions`rg()` and `fd(…, method = “sd”)`and extracted the mean depth of each square. Then, we calculated the total, visible, and cryptic coral diversity (calculated as the total number of species) for each square. Finally, we performed two Bayesian linear regressions predicting coral diversity with depth, rugosity, and fractal dimension including only visible corals, and only cryptic corals, respectively.

Our analysis shows that habitat complexity can influence coral diversity, but patterns differ for visible and cryptic corals (Figure 2). When focusing on visible corals alone, an effect of fractal dimension is detected, while the effect of rugosity is negligible. On the other hand, for cryptic coral diversity, we detect an effect of is rugosity, but not fractal dimension. Depth also has a variable effect on coral diversity: when the reef is shallow, visible corals are more diverse, while cryptic corals are slightly less diverse. This case study exemplifies that structural variables can have nuanced effects on different components of the ecosystem.

**Figure 2.**
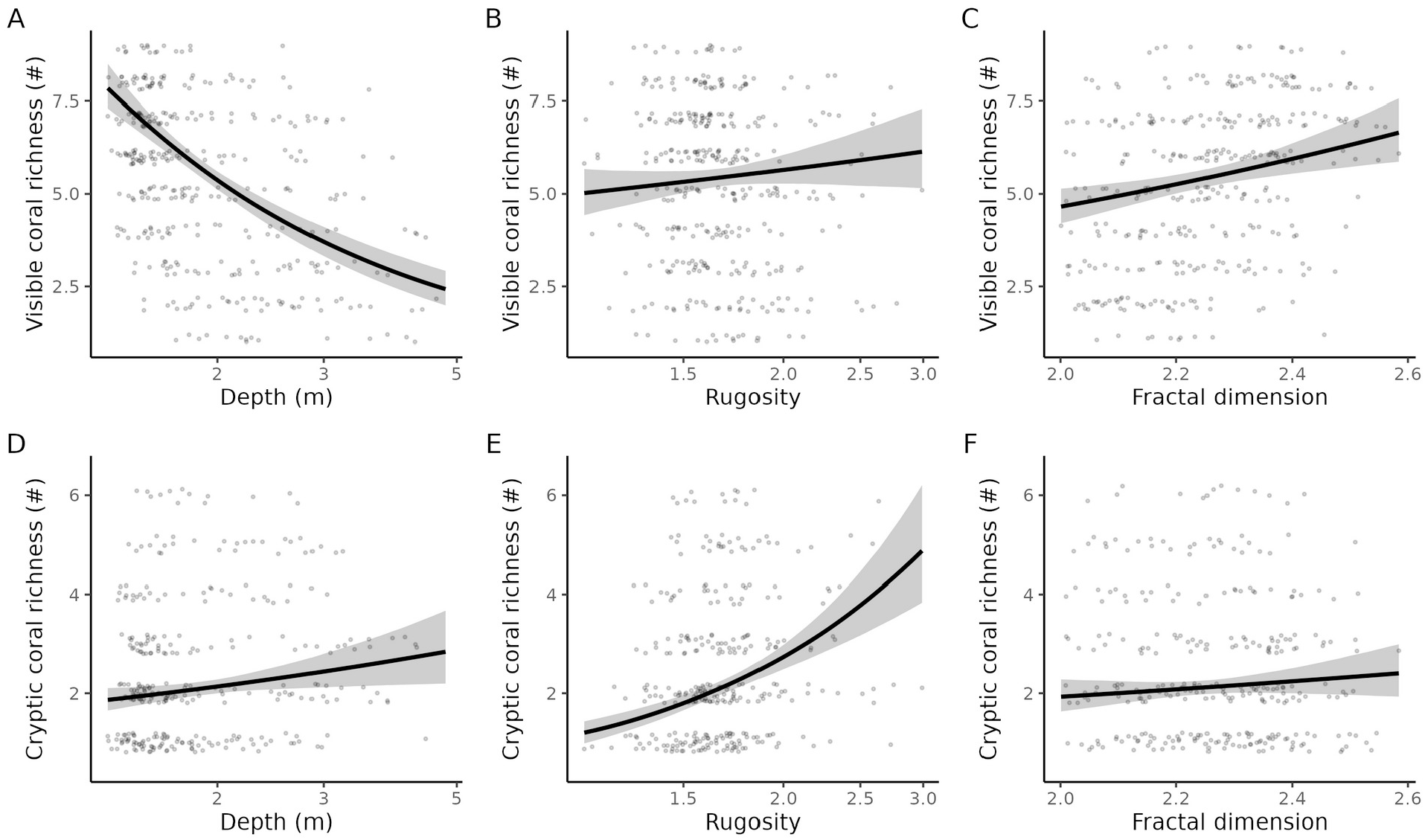
Predicted effects of depth, rugosity, and fractal dimension on visible (A, B, C) and cryptic (D, E, F) coral species richness. Lines show the average predicted values and the shaded area indicates the 95% credible interval of the prediction. Points indicate the raw data and are vertically jittered to increase visibility.

### 3.2 Case study 2: Comparing 3D, 2.5D, and 2D attributes of corals

Many biological and ecological processes are related to the size and shape of the organism, including their life history strategies. However, accurately capturing the 3D morphology of organisms in the field can be difficult, especially when they are numerous and spread over large spatial extents, but their 2D counterparts can be more easily recorded. For example, in coral reefs, 2D top-down images of corals and more recently, large reef photomosaics can be used to trace the outline of corals. These 2D shapes have the potential to be used as a proxy for 3D metrics. For example, 3D colony surface area has been shown to scale consistently with 2D area (House *et al*. 2018). This case study assesses how well 2D and 2.5D metrics (i.e. calculated from a DEM) can approximate 3D shape metrics including rugosity, fractal dimension, and sphericity.

We use `habtools` to quantify 3D attributes of coral colony models generated using laser-scanning, previously collected by Zawada *et al*. (2019) (see supplemental materials). Specifically, we calculate fractal dimension (using the box-counting method), rugosity, and sphericity of each coral mesh. We also convert the 3D mesh objects to DEMs and 2D shapes using `habtools` functions `mesh_to_dem()` and `mesh_to_2d()` and calculate rugosity of each DEM, circularity for 2D shapes, and fractal dimension of both DEMs and 2D shapes (using the standard deviation method and the box-counting method, respectively).

Our results demonstrate positive correlations between equivalent metrics, calculated from various representations of coral colonies (3D mesh, DEM, and 2D shape) (Figure 3). DEM fractal dimensions correlate positively with 3D fractal dimensions and could thus be used as an indicator of 3D fractal dimension. However, it may overestimate 3D fractal dimension when DEM fractal dimension is high. Similarly, 2D fractal dimension correlates with 3D fractal dimension, but there may be consistent bias, depending on the growth form of the corals. For example, 2D fractal dimension is consistently higher than 3D mesh fractal dimension for arborescent corals. Further, 3D mesh rugosity is mostly higher than DEM rugosity, which can be explained by the fact that DEMs do not capture overhangs. For corals with thin branches and plates (e.g. tables and sheet-like corals), estimated DEM rugosity can be much higher than 3D mesh rugosity. This is because a large drop in a DEM will assume vertical walls, which may contribute to a higher estimated surface area. So depending on dominant growth forms in a reef, rugosity could be underestimated or overestimated when relying on DEMs. Finally, circularity can be an indication of sphericity, mostly when an object is similar in each direction (e.g. hemispherical corals). Yet, for table-and sheet-like corals, circularity is a particularly poor indicator of sphericity.

**Figure 3.**
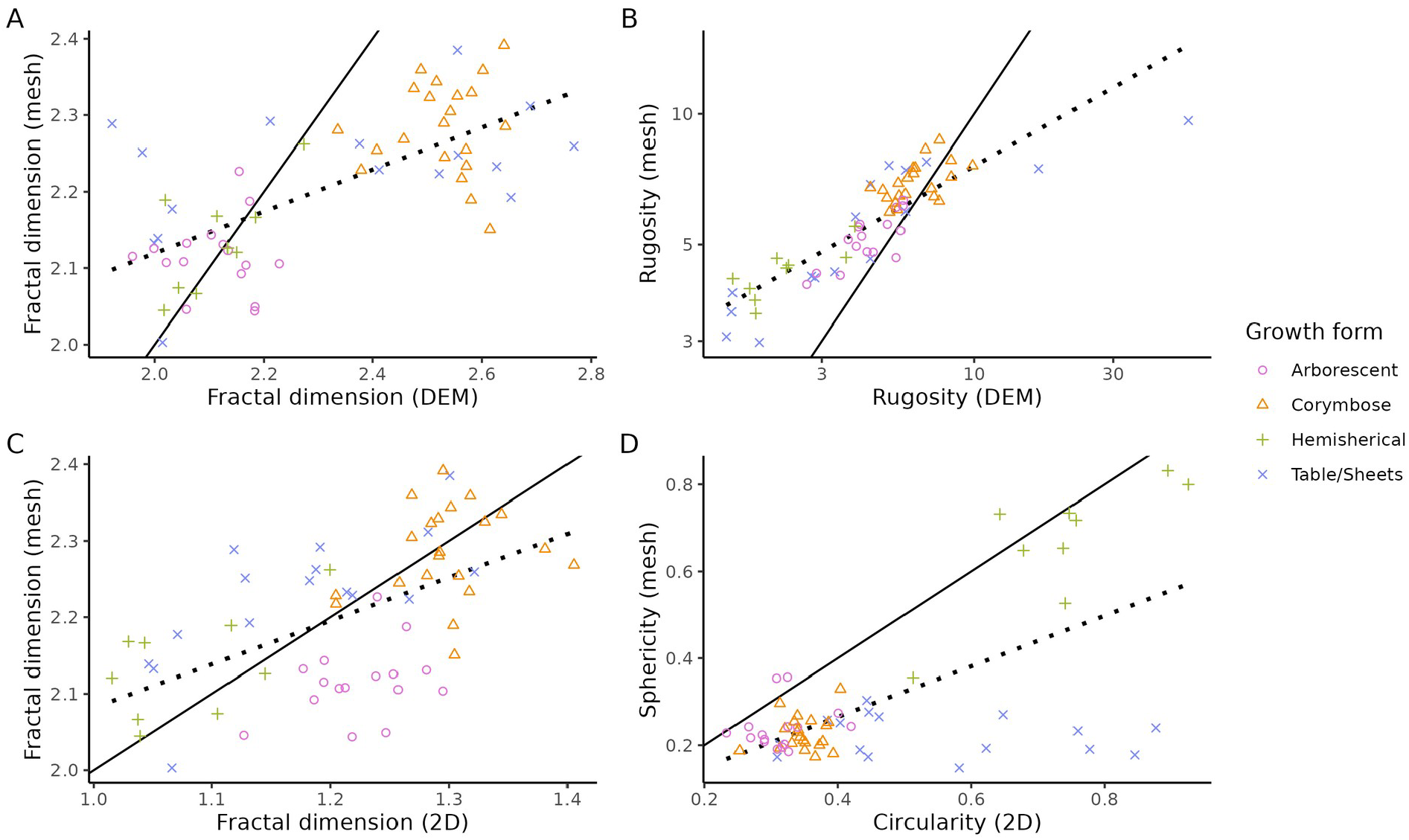
Pairwise comparisons of coral shape metrics. Solid lines indicate the 1:1 line where metrics in both axes are equal. Dotted lines shows the linear regression line for the observed data. Each point represents a coral colony.

## 4. Conclusion

Recent decades have seen extraordinary advances in imaging technologies used to capture the natural world. However, the capability to translate these advances into metrics that can be used in ecological analyses across systems has lagged well behind. The goal of `habtools` is to build and then grow a collection of standard R methods for sampling, simulating, and measuring surfaces and objects. Here, we demonstrated how to best use these methods while highlighting potential pitfalls and biases that are not always obvious to researchers who are new to analyses of surface complexity. We envision that `habtools` will continue to grow as new and faster methods are developed. We welcome new contributions to the package either by direct development of code, or by raising issues and new ideas for functionality via the GitHub repository.

## Supporting information

Case study 1

Case study 2

## Code availability

All code to reproduce the figures included in this manuscript will be available at https://github.com/nschiett/habtools_paper. The DEM and mesh data is available at (TBD). The source code of `habtools` is available at https://github.com/jmadinlab/habtools and can be installed from GitHub for the latest version or directly through CRAN.

## Author contributions

Joshua Madin and Nina Schiettekatte conceived and developed the *habtools* package, lead the analysis, and wrote the first draft of the manuscript; Maria Dornelas, Joshua Madin, Damaris Torres-Pulliza, and Kyle Zawada collected data used as examples in the package and case studies; Mollie Asbury, Guan-Yan Chen, Joshua Madin, Nina Schiettekatte, Damaris Torres-Pulliza, and Kyle Zawada contributed to code development. Mollie Asbury, Guan-Yan Chen, Maria Dornelas, Joshua Madin, Jessica Reichert, Nina Schiettekatte, and Damaris Torres-Pulliza contributed to package testing and package documentation review. All authors contributed critically to the manuscript and gave final approval for publication.

## Acknowledgements

We thank Viviana Brambilla, Garrett Fundakowski, and the many enthusiastic students enrolled in the Marine Biology Graduate Program (MBIO640) at the University of Hawai‘i at Mānoa for package testing and feedback. This work was funded by a National Science Foundation–Natural Environment Research Council Biological Oceanography Grant (1948946) and the European Union (CoralINT, GA 101044975). Views and opinions expressed are however those of the author(s) only and do not necessarily reflect those of the European Union or the European Research Council. Neither the European Union nor the granting authority can be held responsible for them.

## Supplementary material: Fractal dimension

To illustrate how various methods to estimate fractal dimension compare, we applied all four methods to a large number of DEMs including simulated and natural DEMs (extracted from Torres-Pulliza *et al*. (2020). We used two types of algorithms to simulate DEMs: the Diamond-square algorithm (included in `habtools`) and an algorithm using fractional Lévy motions using the `FracSim` R package (Déjean & Cohen 2005). We simulated 190 DEMs for each simulation algorithm: 10 replicates per level of ruggedness. For the reefs, we used the `split_dem()` function to subdivide 8×8m reef sites into 64 smaller squares. All DEMs (natural and simulated) had a fixed number of cells (200×200). We used the same `lvec` across all DEMs for the standard deviation method, height variation method, and the cube counting method (`lvec = L/c(1,2,4,8,16)`, with `L` = the extent of the DEM). For the area method we used a different `lvec` (`lvec = L/c(8,16,32,64,128)`). For the cube counting method and the area method we set `scale = TRUE` inside `fd()`, which rescales the heights so that the height range matches the extent (as done in Fischer & Jucker (2024)).

This method comparison illustrates that fractal dimension methods tend to agree more for simulated fractal surfaces (Supplementary figure 2). Less fractal like surfaces such as natural reefs have more variation, but all methods correlate positively. Moreover, even though we simulated surfaces with a theoretical fractal dimension nearing 3, estimated fractal dimensions tend to be underestimated when surfaces get more complex. Based on our simulated surfaces, the standard deviation method seems to be most accurate (Supplementary figure 2). In summary, when comparing surfaces, users should use a method consistently and be aware of added noise when surfaces are not fractal.

Finally, we demonstrate how resolution and height range affect the fractal dimension estimates using various methods (Supplementary figure 4). First, we altered the resolution of natural reef DEMs and estimated the fractal dimension values. We found that the resolution affects the fractal dimension estimate when using the height variation method and the standard deviation method (Supplementary figure 4B), while resolution does not affect the fractal dimension estimate when using the cube counting or area method. Second, we assessed the effect of height on fractal dimension and found that it heavily affects the estimates when using the cube counting and area methods (Supplementary figure 4A). The height variation and standard deviation methods are invariant to the height range.

## Supplementary figures

**Supplementary figure 1.**
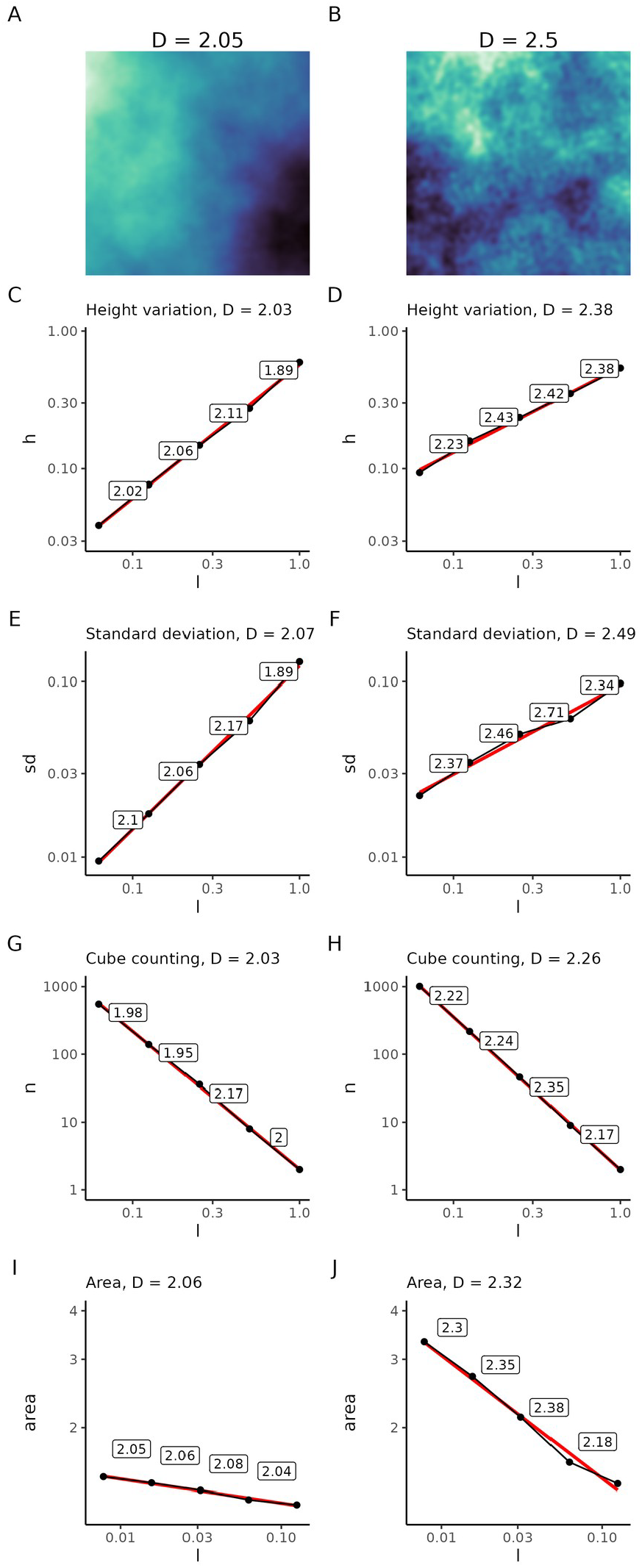
Examples of fractal dimension calculation using multiple methods for a DEM with low complexity (A) and a DEM with high complexity (B). Plots show the diagnostic plots for both DEMs using the height variation method (C, D), the standard deviation method (E, F), the cube counting method (G, H), and the area method (I,J). These plots are similar to the plots produced when `diagnose = TRUE` inside the `fd()` function. The red lines indicates the overall regression lines and the labels show the computed fractal dimensions between each scale transition.

**Supplementary figure 2.**
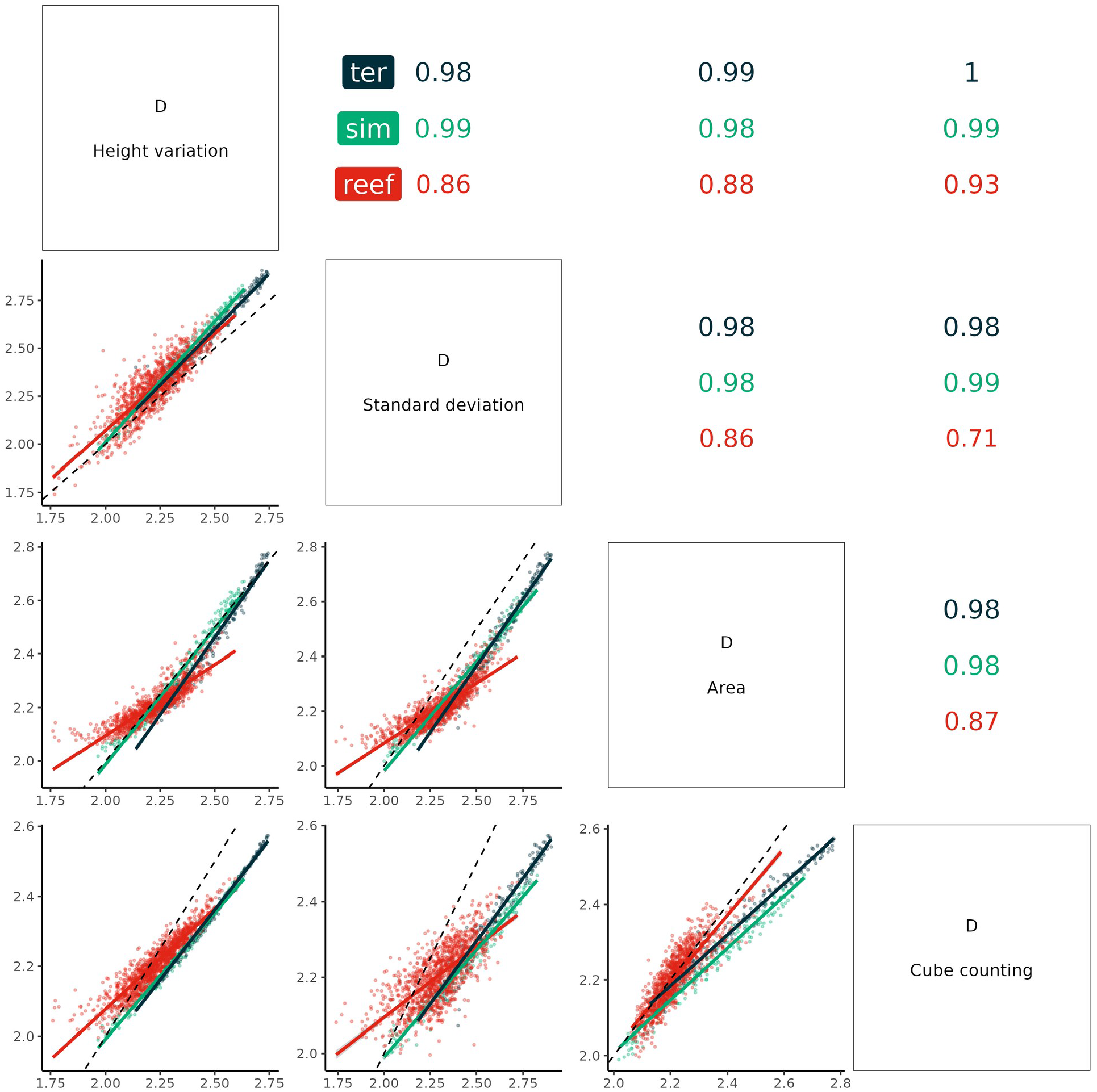
Correlation plot comparing estimated fractal dimensions (D) using various methods including the height variation method, the standard deviation method, the area method, and the cube counting method. DEMs for this comparison included 190 simulated surfaces using the diamond-square algorithm (included in habtools through the function `sim_dem()`, black), 190 simulated surfaces using fractional Lévy motions (sim, green), and 1216 reef patches (reef, red). Numbers indicate the correlation coefficients. Unity line of slope 1, intercept 0 is given for comparison (black dashed).

**Supplementary figure 3.**
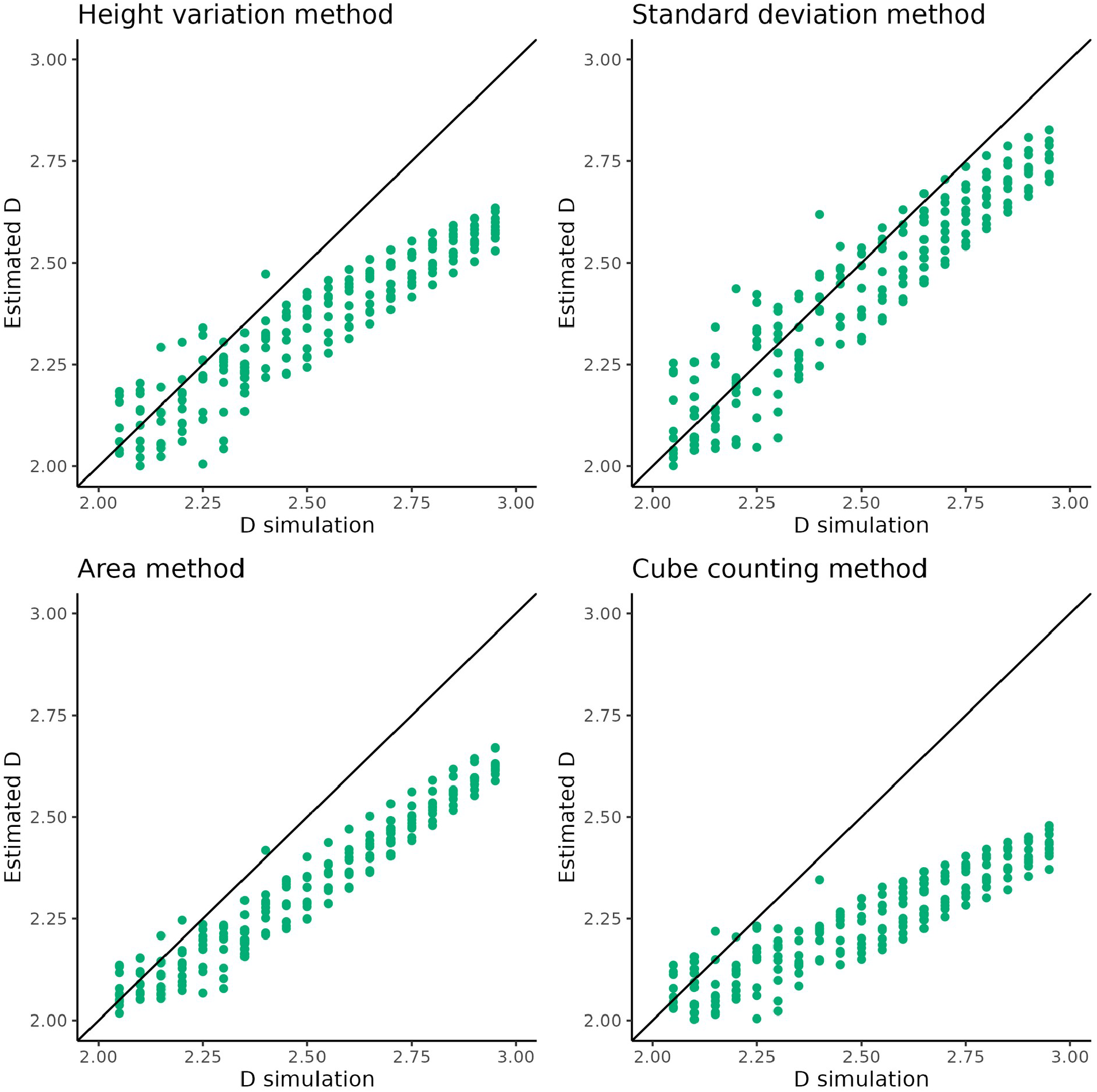
Comparison of the theoretical fractal dimension specified for the simulation of DEMs (using fractional Lévy motions) and estimated fractal dimensions using various methods. The black line indicates the regression line a slope of 1.

**Supplementary figure 4.**
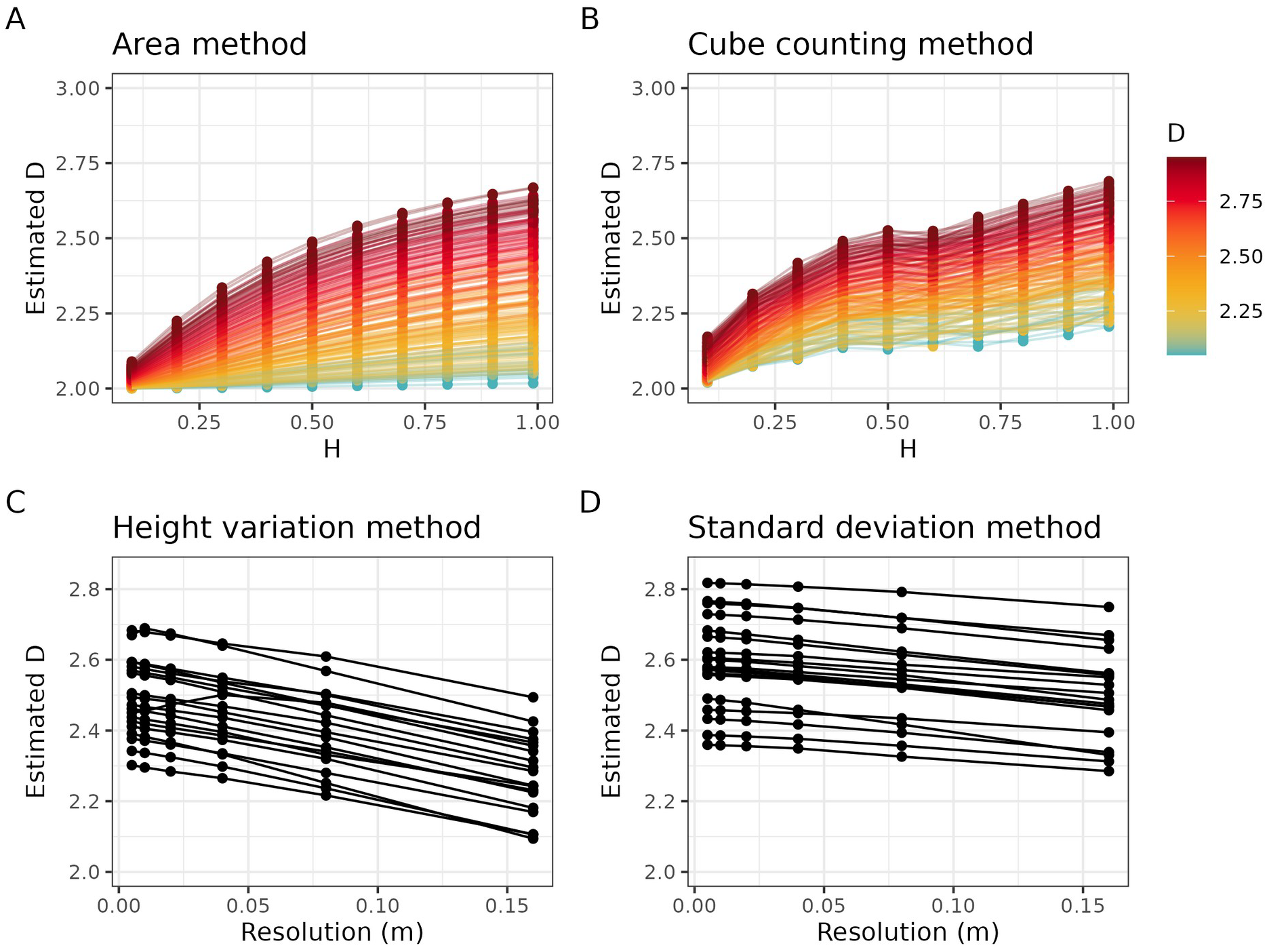
Potential sources of variation affecting fractal dimension estimation. The estimated fractal dimension (D) increases with increasing height range when using the area method (A) or the cube counting method (B). The color scale indicates the D used for the simulation using fractional Lévy motions. Estimated D decreases with increased resolution when using the height variation method (C) or the standard deviation method (D). Lines connect the estimated values for the same DEM across varying height range (H) and resolution. To assess fractal dimension calculations across resolutions, we used 8×8 natural reef DEMs from Torres-Pulliza *et al*. (2020).

