## Supplementary material for "habtools: an R package to calculate 3D metrics for surfaces and objects": Case study 1

### Case study 1: The effect of rugosity and fractal dimension on coral diversity

#### Summary

Both rugosity and fractal dimension have been shown to explain variation in coral species richness, expressed as the number of unique species observed (Torres-Pulliza *et al.* 2020). Corals are ecosystem engineers so the link between structure and diversity could be related to the fact that corals provide structure. For this case study, we investigate how the relationship between complexity metrics and coral diversity varies when considering visible corals versus cryptic corals. We use the reef structure and coral diversity data examined in Torres-Pulliza *et al.* (2020). Using Structure-from-motion, a digital elevation model (DEM) of a large reef patch (~562m$^2$) near Lizard Island, Australia was constructed. All corals were annotated and identified in the field, resulting in a shape file of annotated dots that can be spatially matched with the DEM. The annotated corals included both visible corals (i.e. seen from a top-down view) and cryptic corals (i.e. hidden under a ledge or in a crevice). Here, we illustrate how `habtools` can be used to link structural complexity and biodiversity (see supplementary materials for a detailed description of the methods and code). Specifically, we compare how structural complexity metrics (rugosity and fractal dimension) correlate with visible corals and cryptic corals, respectively.

We use the `dem_sample()` function to sample 500 square DEMs with an side of 1m out of the larger reef patch. For each smaller square (1m$^2$), we quantified rugosity and fractal dimension using functions`rg()` and `fd(..., method = "sd")`and extracted the mean depth of each square. Then, we calculated the total, visible, and cryptic coral diversity (calculated as the total number of species) for each square. Finally, we performed two Bayesian linear regressions predicting coral diversity with depth, rugosity, and fractal dimension including only visible corals, and only cryptic corals, respectively.

Our analysis shows that habitat complexity can influence coral diversity, but patterns differ for visible and cryptic corals (Figure 2). When focusing on visible corals alone, an effect of fractal dimension is detected, while the effect of rugosity is negligible. On the other hand, for cryptic coral diversity, we detect an effect of is rugosity, but not fractal dimension. Depth also has a variable effect on coral diversity: when the reef is shallow, visible corals are more diverse, while cryptic corals are slightly less diverse. This case study exemplifies that structural variables can have nuanced effects on different components of the ecosystem.

Below, we illustrate how we used `habtools` functions to perform this analysis.

#### Code

##### Load R packages

```
library(readr)
library(tidyr)
library(dplyr)
library(sf)
library(raster)
library(ggplot2)
library(habtools)
library(patchwork)
library(brms)
```

##### Load reef

We start by loading the data, including the DEM of the reef and a shape file of coral species. The coral data includes information about the visibility of the corals. We classify corals that were hidden under a structure, inside a gap, or on a vertical wall as cryptic.

```r
# Reef DEM
reef <- raster::raster("data/megaplot/trimodal_cropped.tif")

# Shape file with annotated corals
corals <- st_read("data/megaplot/trimodal_ann_aligned_cleaned.shp") %>%
  dplyr::filter(!Visibility == "0") %>%
  dplyr::select(OBJECTID, X, Y, Species, Visibility) %>%
  dplyr::mutate(cryptic = case_when(
    Visibility %in% c("under", "gap", "vertical") ~ "cryptic",
    TRUE ~ "visible"))
```

```
## Reading layer ‘trimodal_ann_aligned_cleaned’ from data source
##   ‘/home/nina/Documents/work/himb_projects/habtools_paper/data/megaplot/trimodal_ann_aligned_cleaned
##   using driver ‘ESRI Shapefile’
## Simple feature collection with 9264 features and 31 fields
## Geometry type: POINT
## Dimension:     XY
## Bounding box:  xmin: -125.7191 ymin: -49.80476 xmax: -72.49465 ymax: -19.42645
## Projected CRS: Transverse_Mercator
```

```r
raster::plot(reef)
```

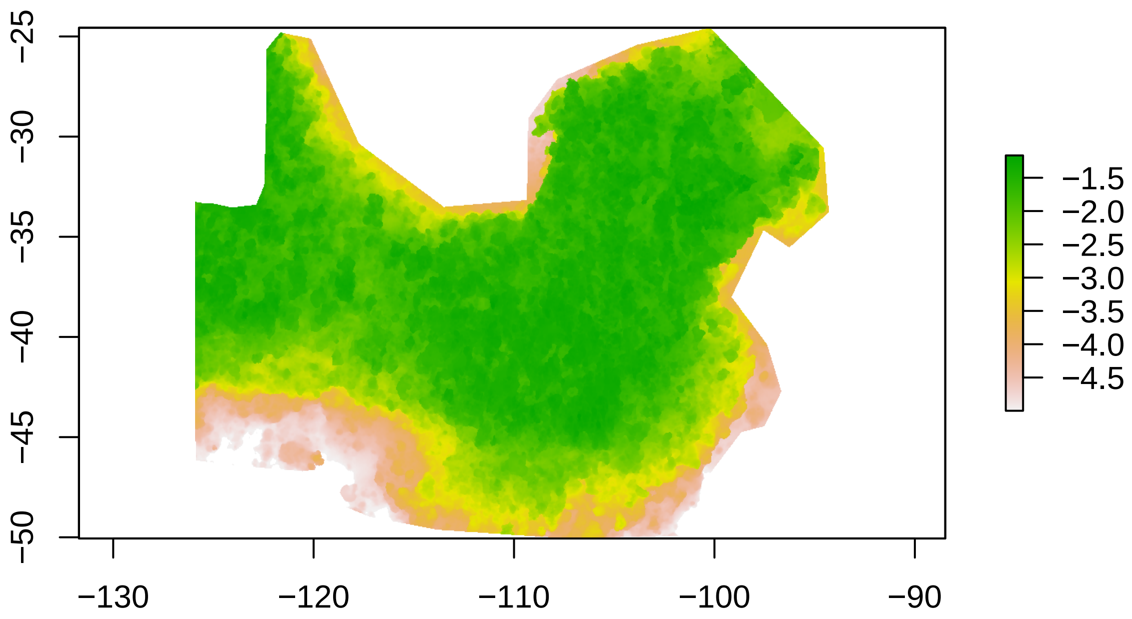

##### Sample a single DEM

Using `habtools` function `dem_sample()`, we can crop a random square (1x1m) out of the reef for further analysis. By setting `allow_NA` to 0, we avoid any cells with NAs. When setting plot = TRUE, we can visualize the sample location. Then, we can use functions `fd()`, `rg()`, and `fd()` to calculate fractal dimension, rugosity, and height range, respectively. In this case study, we chose to use the "sd" method for calculating fractal dimension.

```
dem <- dem_sample(reef, L = 1, plot = TRUE, allow_NA = 0)
```

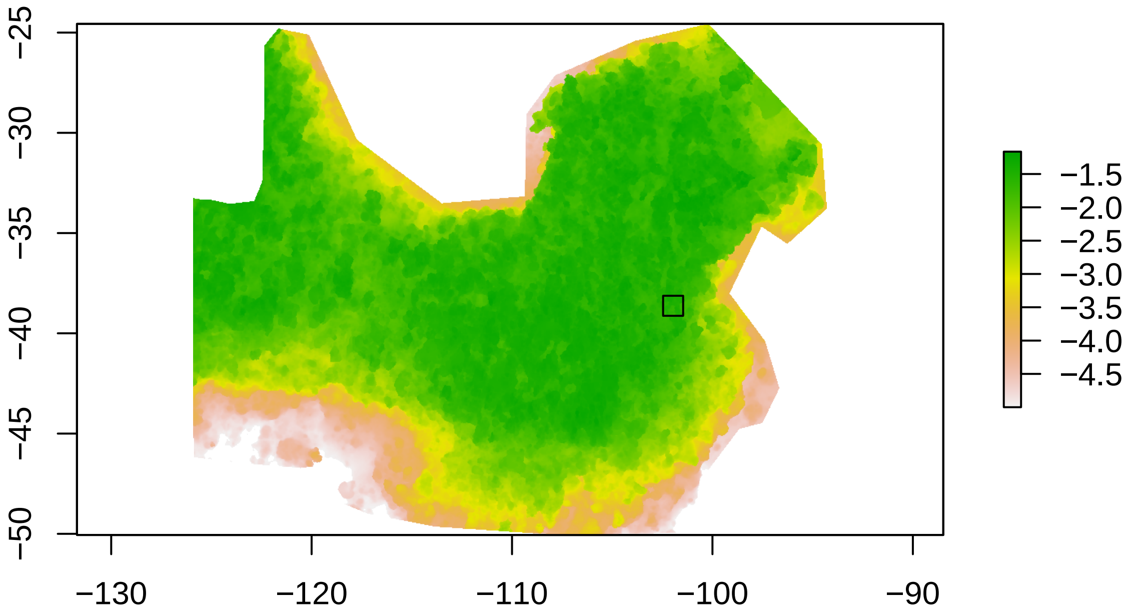


```
plot(dem)
```

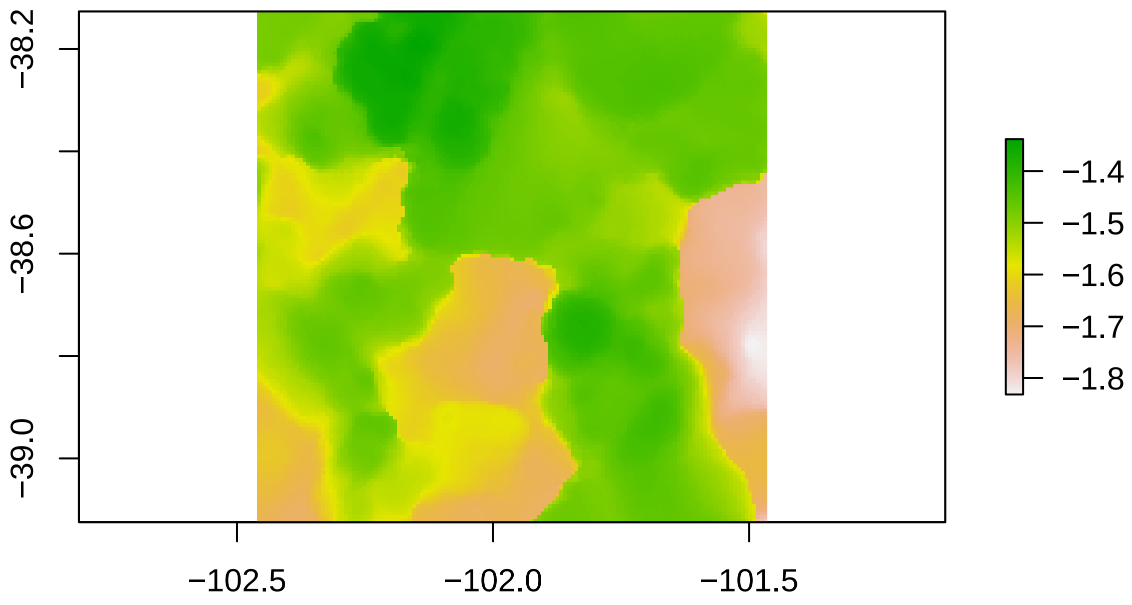


```r
fd(dem, lvec = c(1, 0.5, 0.25, 0.125), method = "sd")
```

```
## [1] 2.399561
```

```r
rg(dem)
```

```
## [1] 1.674574
```

```r
hr(dem)
```

```
## [1] 0.4936714
```

We can now match the location of the sampled DEM with the coral shape file and extract the coral species richness.

```r
# construct a shape file of the bounding box of the DEM
box <- st_as_sfc(st_bbox(dem), crs = crs(dem))
# Filter out the corals that are in the area of interest
corals_box <- st_intersection(corals, box)

# species richness
length(unique(corals_box$Species))
```

```
## [1] 13
```

```r
# species richness cryptic vs visible
corals_box %>%
  as.data.frame() %>%
  dplyr::group_by(cryptic) %>%
  dplyr::summarise(abu = n(), nspec = length(unique(Species)))
```

```
## # A tibble: 2 x 3
##   cryptic   abu nspec
##   <chr>   <int> <int>
## 1 cryptic     1     1
## 2 visible    16    12
```

###### Sample multiple squares

To assess the relationship between structural complexity and coral diversity, we want to get a wide spread of reef patches with variable rugosity and fractal dimension values. Therefore, we sample multiple squares and calculate complexity and coral richness metrics. In the code below, we sample 10 squares (n = 10) in a for loop. At each iteration, we replace the center cell of the sampled square by NA before sampling the reef again to limit overlap between samples to less than 50%.

```r
# create list of sampled DEMs
dems <- list()
n <- 10
L <- 1

plot(reef)
for(i in 1:n) {
  dem <- dem_sample(data = reef, L = L,
                    allow_NA = FALSE, plot = FALSE, max_iter = 200)
  dems[[i]] <- dem
  mid <- habtools::mid_find(dem)
  x0 <- mid$x_mid
  y0 <- mid$y_mid
  rect(x0 - (L/2), y0 - (L/2), x0 + (L/2), y0 + (L/2))
  # replace center of square by NA in reef
  icell <- cellFromXY(reef, mid)
  reef[icell] <- NA
}
```

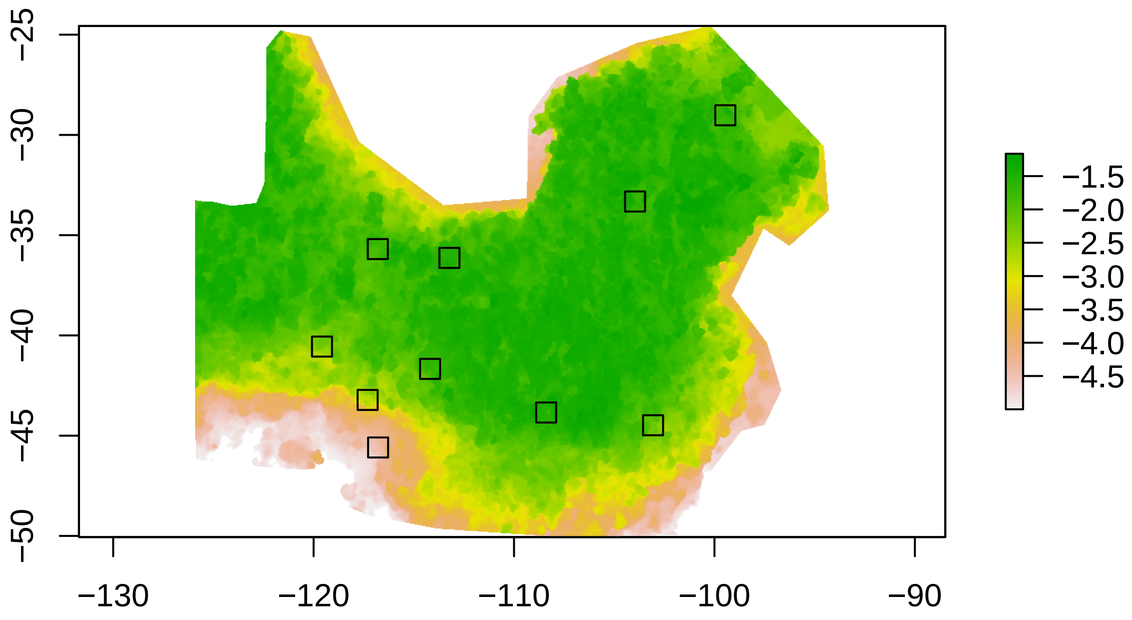


Then, we can estimate complexity metrics for each of the sampled squares.

```
# rugosity
r <- sapply(dems, rg)
# height range
h <- sapply(dems, hr)
# Fractal dimension
d <- sapply(dems, fd, method = "sd", lvec = c(1, 0.5, 0.25, 0.125))
# Mean depth of square
z <- sapply(dems, z)
# mid points coordinates of squares
mids <- lapply(dems, mid_find) %>% bind_rows()

# combine into data frame
data_rdh <- data.frame(id = 1:length(dems), r = r, d = d, h = h, z = z) %>%
  cbind(mids)
```

We can also extract coral species richness for each square.

```
data_corals <- lapply(1:length(dems), function(i){
  # bounding box
  e <- st_as_sfc(st_bbox(dems[[i]]),
                 crs = crs(dems[[i]]))
  # Filter out the corals that are in the area of interest
  asub <- st_intersection(corals, e)
  if(nrow(asub)>0) {
```

```
    asub$id <- i
  }
  asub
}) %>% plyr::ldply() %>%
  dplyr::group_by(id) %>%
  dplyr::mutate(abu_all = n(),
                nspec_all = length(unique(Species))) %>%
  dplyr::group_by(id, cryptic,abu_all, nspec_all) %>%
  dplyr::summarise(abu = n(),
                   nspec = length(unique(Species))) %>%
  pivot_wider(names_from = cryptic, values_from = c(abu, nspec), values_fill = 0)
```

Because we added an id code for each square, we can combine both data sets.

```
data <- left_join(data_rdh, data_corals)
head(data)
```

```
##   id        r        d         h         z      x_mid     y_mid abu_all nspec_all
## 1  1 1.353275 2.252551 0.4758372 -1.766651 -116.79636 -35.69279      10        10
## 2  2 1.749262 2.302806 0.5810051 -1.704342  -99.45857 -29.01940       8         4
## 3  3 1.791829 2.195944 0.7706857 -2.116449 -103.06185 -44.48055       9         6
## 4  4 1.593354 2.152030 0.6443267 -1.744015 -114.18886 -41.66976       6         5
## 5  5 1.364043 1.950811 0.8526945 -4.424312 -116.78160 -45.58979       2         2
## 6  6 2.347386 1.903141 1.4270554 -2.890862 -117.30531 -43.21695       3         3
##   abu_cryptic abu_visible nspec_cryptic nspec_visible
## 1           2           8             2             8
## 2           1           7             1             4
## 3           4           5             4             2
## 4           1           5             1             4
## 5           1           1             1             1
## 6           0           3             0             3
```

##### Data analysis

By changing `n = 10` to ‘`n = 500` in the above code, we can collect the replicates needed for our analysis. We ran this code 500 squares so for the purpose of avoiding long computation time, one can load this data set. For this analysis, we filter out outliers (i.e. rugosity higher than 3 and fractal dimension lower than 2) and consider only squares that have at least one coral in them.

```
data <- read_csv("output/mega_data.csv") %>%
  filter(r<=3, nspec_all > 0, d>2) %>%
  mutate(z = abs(z))
```

We can visualize the data and potential relationships.

```
ggplot(data) +
  geom_point(aes(x = z, y = nspec_visible), alpha = 0.5) +
  labs(x = "Depth (m)", y = "Visible coral richness (#)") +

  ggplot(data) +
  geom_point(aes(x = r, y = nspec_visible), alpha = 0.5) +
```

```
  labs(x = "Rugosity", y = "Visible coral richness (#)") +
  scale_x_log10() +

  ggplot(data) +
  geom_point(aes(x = d, y = nspec_visible), alpha = 0.5) +
  labs(x = "Fractal dimension", y = "Visible coral richness (#)") +


  ggplot(data) +
  geom_point(aes(x = z, y = nspec_cryptic), alpha = 0.5) +
  labs(x = "Depth (m)", y = "Cryptic coral richness (#)") +


  ggplot(data) +
  geom_point(aes(x = r, y = nspec_cryptic), alpha = 0.5) +
  labs(x = "Rugosity", y = "Cryptic coral richness (#)") +
  scale_x_log10() +

  ggplot(data) +
  geom_point(aes(x = d, y = nspec_cryptic), alpha = 0.5) +
  labs(x = "Fractal dimension", y = "Cryptic coral richness (#)") &

  theme_classic()
```

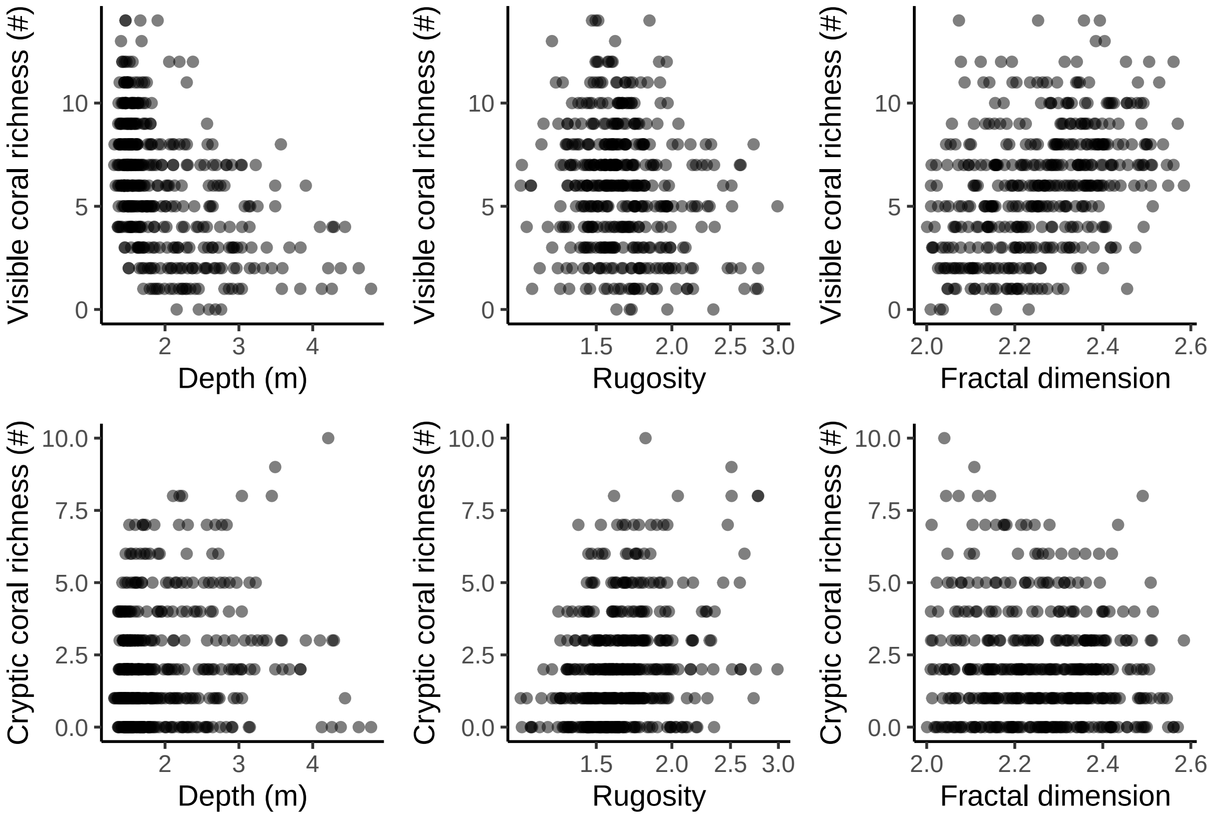


To asses the effect of depth (z), rugosity (r), and fractal dimension (d) on coral diversity (y), we perform a Bayesian linear regression with a poisson distribution and a log-link function:

$$mu = z + log(r) + dy \sim poisson(exp(mu)).$$

We perform this regression for both visible corals and cryptic corals using the R package `brms` (Burkner 2017).

```
fit_visible <- brm(nspec_visible ~ log(r) + z + d,
                   family = "poisson",
                   data = data, backend = "cmdstanr",
                   silent = 2)

summary(fit_visible)
```

```
##  Family: poisson
##   Links: mu = log
## Formula: nspec_visible ~ log(r) + z + d
##    Data: data (Number of observations: 432)
##   Draws: 4 chains, each with iter = 2000; warmup = 1000; thin = 1;
##          total post-warmup draws = 4000
##
## Regression Coefficients:
##           Estimate Est.Error l-95% CI u-95% CI Rhat Bulk_ESS Tail_ESS
## Intercept     0.85      0.48    -0.09     1.77 1.00     3070     2353
## logr          0.13      0.15    -0.16     0.41 1.00     3807     2924
## z            -0.38      0.05    -0.48    -0.30 1.00     3302     2740
## d             0.70      0.18     0.35     1.06 1.00     3219     2660
##
## Draws were sampled using sample(hmc). For each parameter, Bulk_ESS
## and Tail_ESS are effective sample size measures, and Rhat is the potential
## scale reduction factor on split chains (at convergence, Rhat = 1).
```

The regression for visible coral richness shows a negative effect of depth (-0.38, 95%CI: -0.47;-0.30) and a positive effect of fractal dimension (0.70, 0.34;1.07). The effect of rugosity was low and the 95% credible interval overlapped with zero.

```
fit_cryptic <- brm(nspec_cryptic ~ log(r) + z + d,
                   family = "poisson",
                   data = data, backend = "cmdstanr",
                   silent = 2)

summary(fit_cryptic)
```

```
##  Family: poisson
##   Links: mu = log
## Formula: nspec_cryptic ~ log(r) + z + d
##    Data: data (Number of observations: 432)
##   Draws: 4 chains, each with iter = 2000; warmup = 1000; thin = 1;
##          total post-warmup draws = 4000
##
## Regression Coefficients:
##           Estimate Est.Error l-95% CI u-95% CI Rhat Bulk_ESS Tail_ESS
## Intercept    -1.00      0.81    -2.60     0.61 1.00     2840     2542
## logr          1.48      0.21     1.07     1.91 1.00     3861     3216
```

```
## z              0.12      0.06     0.01     0.23 1.00     3239     2871
## d              0.32      0.31    -0.30     0.94 1.00     2973     2616
##
## Draws were sampled using sample(hmc). For each parameter, Bulk_ESS
## and Tail_ESS are effective sample size measures, and Rhat is the potential
## scale reduction factor on split chains (at convergence, Rhat = 1).
```

The regression for cryptic coral richness shows a slight positive effect of depth (0.12, 95%CI: 0.01;0.23) and a strong positive effect of rugosity (1.48, 1.07;1.90). The effect of fractal dimension was low and the 95% credible interval overlapped with zero.

We can visualize the results by plotting the partial effects.

```r
me_visible <- conditional_effects(fit_visible)
me_cryptic <- conditional_effects(fit_cryptic)

ggplot(me_visible[[2]]) +
  geom_ribbon(aes(x = z, ymin = lower__, ymax = upper__), alpha = 0.3) +
  geom_smooth(aes(x = z, y = estimate__), color = "black") +
  labs(x = "Depth (m)", y = "Visible coral richness (#)") +
  scale_y_continuous(limits = c(1,9)) +

  ggplot(me_visible[[1]]) +
  geom_ribbon(aes(x = r, ymin = lower__, ymax = upper__), alpha = 0.3) +
  geom_smooth(aes(x = r, y = estimate__), color = "black") +
  labs(x = "Rugosity", y = "Visible coral richness (#)") +
  scale_y_continuous(limits = c(1,9)) +

  ggplot(me_visible[[3]]) +
  geom_ribbon(aes(x = d, ymin = lower__, ymax = upper__), alpha = 0.3) +
  geom_smooth(aes(x = d, y = estimate__), color = "black") +
  labs(x = "Fractal dimension", y = "Visible coral richness (#)") +
  scale_y_continuous(limits = c(1,9)) +

  ggplot(me_cryptic[[2]]) +
  geom_ribbon(aes(x = z, ymin = lower__, ymax = upper__), alpha = 0.3) +
  geom_smooth(aes(x = z, y = estimate__), color = "black")+
  labs(x = "Depth (m)", y = "Cryptic coral richness (#)") +
  scale_y_continuous(limits = c(0.5,6.5)) +

  ggplot(me_cryptic[[1]]) +
  geom_ribbon(aes(x = r, ymin = lower__, ymax = upper__), alpha = 0.3) +
  geom_smooth(aes(x = r, y = estimate__), color = "black") +
  labs(x = "Rugosity", y = "Cryptic coral richness (#)") +
  scale_y_continuous(limits = c(0.5,6.5)) +

  ggplot(me_cryptic[[3]]) +
  geom_ribbon(aes(x = d, ymin = lower__, ymax = upper__), alpha = 0.3) +
  geom_smooth(aes(x = d, y = estimate__), color = "black") +
  labs(x = "Fractal dimension", y = "Cryptic coral richness (#)") +
  scale_y_continuous(limits = c(0.5,6.5)) +

  plot_layout(ncol = 3) + plot_annotation(tag_levels = "A") &
  theme_classic()
```

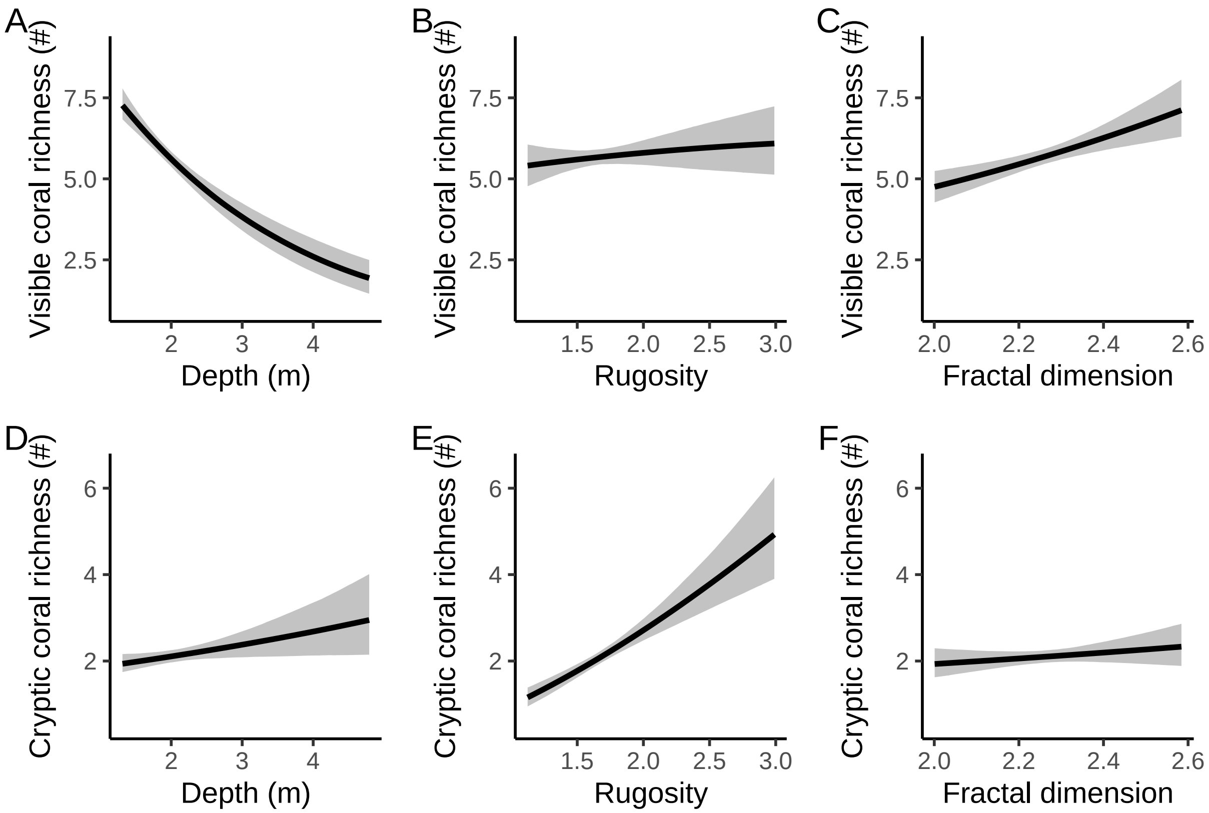
