## Supplementary material for "habtools: an R package to calculate 3D metrics for surfaces and objects": Case study 2

Below, we provide an example of the workflow for a single coral colony, then apply the same approach for a list of coral models, and finally visualize pairwise comparisons of metrics.

### Code

#### Mesh conversion

We use a collection of 3D coral models capturing the morphological diversity of reef building corals (Zawada *et al.* 2019). Models in `ply` format were loaded in R using the `Rvcg` package (Schlager 2017). We then use the `habtools` function `mesh_to_2d()` to convert the 3D meshes into the 2D planar projected shapes (xy coordinates) and `mesh_to_dem()` to convert 3D meshes into DEMs. To exemplify the steps, we first focus on a single massive coral colony.

```
# load mesh
mesh <- Rvcg::vcgPlyRead("data/meshes/zawada/BP10MaPsppLZPROWtFabDATColony.ply")
# Inspect resolution
Rvcg::vcgMeshres(mesh)[1]
## $res
## [1] 0.5092938
```

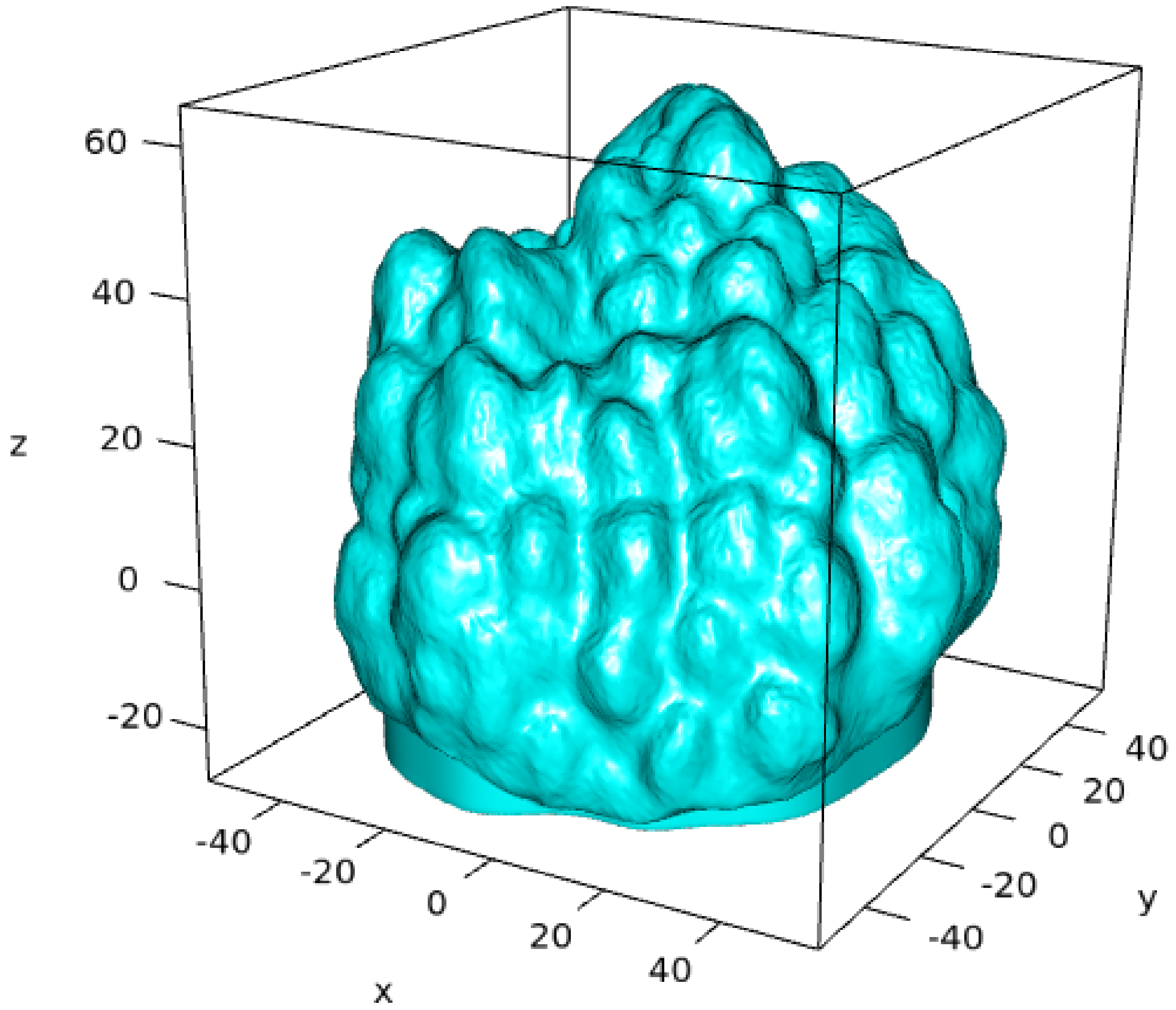


```
# create planar outlines xy coordinates from mesh
shape <- habtools::mesh_to_2d(mesh)
plot(shape, type = "l")
```

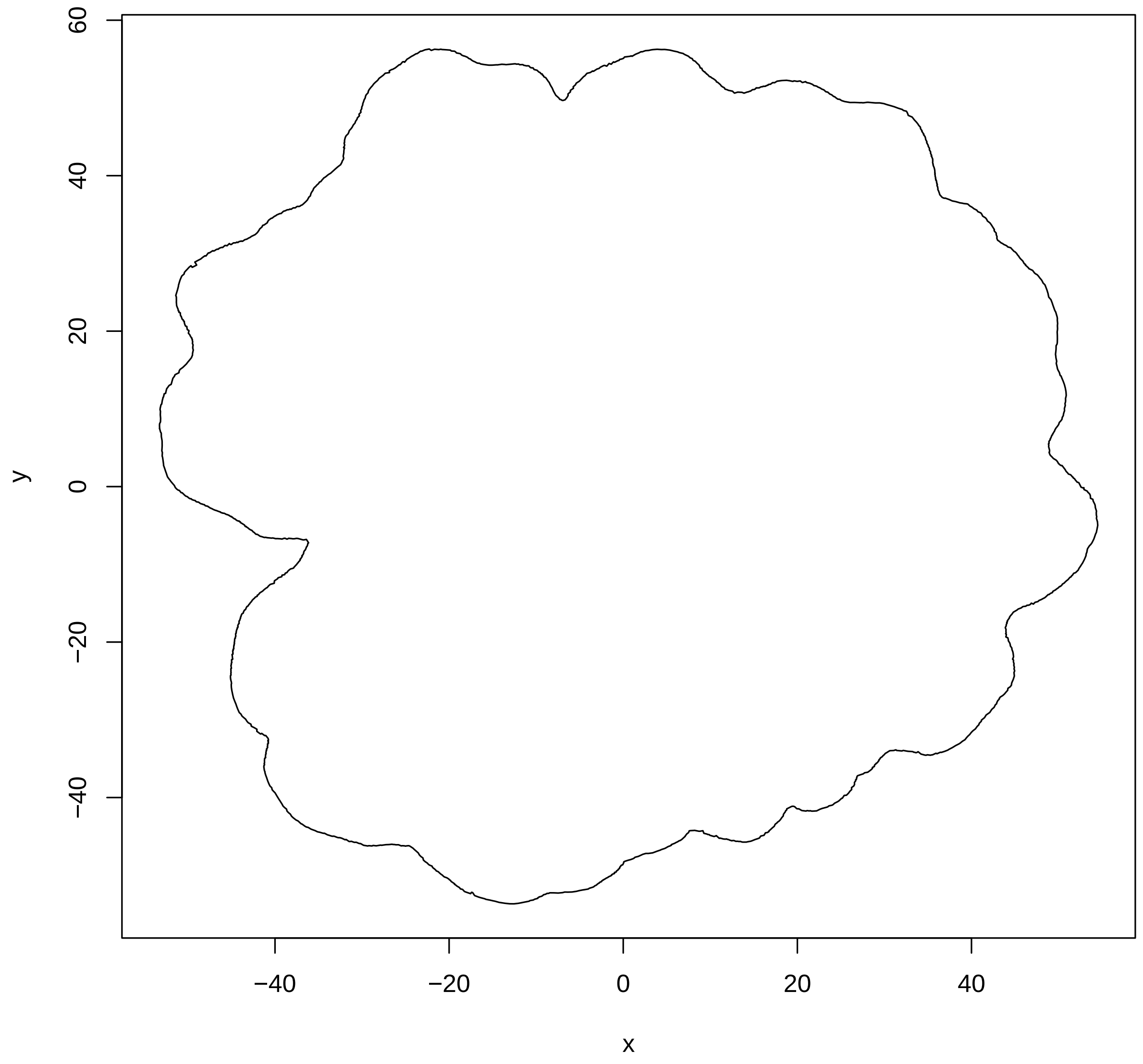


```r
# create DEM from mesh
dem <- habtools::mesh_to_dem(mesh, res = 1, fill = FALSE)
raster::plot(dem)
```

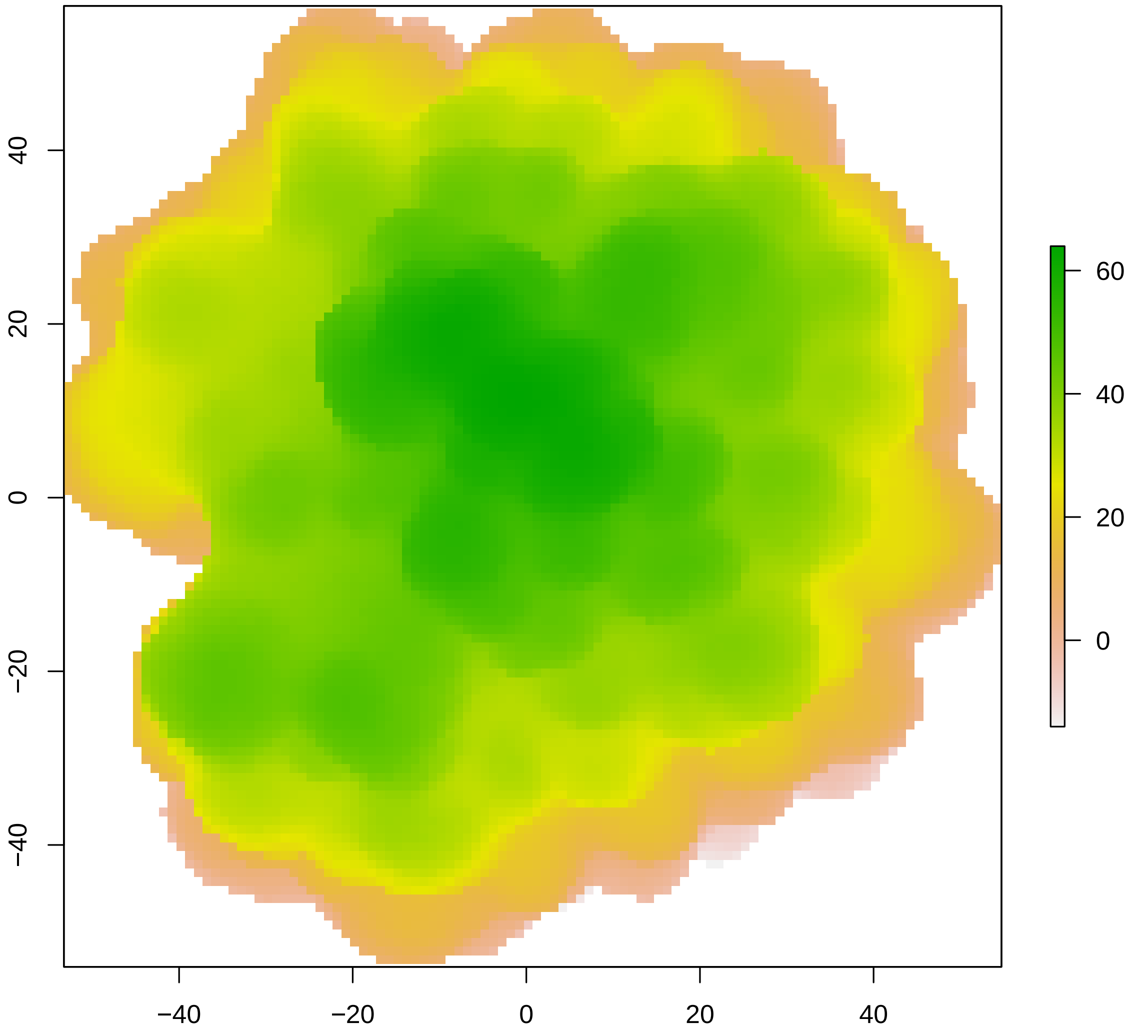


#### Calculate metrics

**Fractal dimension** We calculate fractal dimension of the coral colony using the mesh itself, the DEM, and the 2D shape. To estimate fractal dimension, we opt for scales ranging between 5 and 50, ensuring that the lowest scale exceeds the resolution of our object. We chose the highest value in lvec to be 50 to avoid including the scale transition between highest scales. For the fractal dimension calculation of the DEM, we first need to sample a square from the original DEM because the methods require the width and height of the DEM to be equal. We sample five random squares from the larger DEM. We specified the side length to be 100 (i.e. the largest possible square while ensuring the ability to be subdivided by each scale in lvec).

```
# lvec ranging between 5 and 50
lvec <- 50/c(10,6,3,2,1)
lvec
```

```
## [1]  5.000000  8.333333 16.666667 25.000000 50.000000
# calculate fractal dimension of mesh
fd(mesh, lvec = lvec, method = "cubes")
## [1] 2.126345
# calculate fractal dimension of dem
fd(shape, lvec = lvec, method = "boxes")
## [1] 1.076643
# sample random square dems with fixed size from our DEM
dem_list <- lapply(1:5, function(i){
         dem_sample(dem, 100, allow_NA = 0.5, plot = FALSE)})
# calculate fractal dimension for each sampled dem
fd_dems <- sapply(dem_list, fd, lvec = lvec, method = "sd", parallel = T)
fd_dems
## [1] 2.132622 2.137649 2.130295 2.142924 2.137199
# average fractal dimension of the dem
mean(fd_dems)
## [1] 2.136138
```

#### Rugosity

We calculate rugosity (i.e. the ratio between surface area and planar area) for both the original mesh and the DEM, using the same resolution (1mm).

```
rg(mesh, L0 = 1)
## [1] 4.168434
rg(dem, L0 = 1)
## [1] 2.1916
```

The rugosity calculated from the mesh is much higher (almost double) than the rugosity calculated from the DEM. This is to be expected as the DEM ignores any overhangs. This is why DEMs are sometimes called 2.5D.

#### Sphericity and circularity

A perfect sphere would have a sphericity of 1 and if we transform a sphere into a 2D shape, we would get a perfect circle with a circularity of 1. If 3D shapes are similar in each direction, we could hypothesize that the sphericity of 3D shapes and the circularity of the associated 2D outlines are strongly correlated. We estimate sphericity of the mesh and circularity of the 2D shape.

```
sphericity(mesh)
## [1] 0.7845665
circularity(shape)
## [1] 0.8360726
```

#### Metrics for multiple coral colonies

We can use the above-described approach to estimate metrics for a list of coral colonies of various sizes. For our case study, we only consider colonies that are larger than 100mm in each dimension. CAUTION: this code is computationally expensive and may take hours to run.

```
# get filenames
files <- dir("data/meshes/zawada/")

# subset meshes that are bigger than 100mm in each dimension
lcheck <- sapply(files, function(x){
  print(which(x == files))
  # load meshes
  mesh <- Rvcg::vcgPlyRead(paste0("data/meshes/zawada/", x))
  dt <- t(mesh$vb)
  floor(min(c(
    abs(max(dt[,1]) - min(dt[,1])),
    abs(max(dt[,2]) - min(dt[,2])),
    abs(max(dt[,3]) - min(dt[,3])))))
})

files_sub <- files[which(lcheck >100)]

# apply calculations for each mesh
out <- lapply(files_sub, function(x){
  print(which(x == files_sub))
  # load meshes
  mesh <- Rvcg::vcgPlyRead(paste0("data/meshes/zawada/", x))

  # create planar outlines xy coordinates from meshes
  shape <- habtools::mesh_to_2d(mesh)

  # transform to square dem
  dem <- habtools::mesh_to_dem(mesh, res = 1, fill = FALSE)

  dem_list <- lapply(1:5, function(i){
    print(i)
    dem_sample(dem, 100, allow_NA = 0.5, plot = F, max_iter = 500)})

  d_dem <- sapply(dem_list, fd, parallel = T, method = "sd",
                  lvec = lvec)

  # Create dataframe with scanID
  data <- data.frame(ScanID = gsub( "\\..*$", "", x))

  # fd
  data$D3_dem <- mean(d_dem)
  # 3D fractal dimension
  data$D3_mes <- fd(mesh, lvec = lvec,  method="cubes")
  # 2D fractal dimension
  data$D2_shp <- fd(shape, method = "boxes", lvec = lvec)

  # mesh metrics
  data$R_mes <- rg(mesh, L0 = 1)
  data$spher_mes <- habtools::sphericity(mesh)

  # shape metrics
  data$circ_shp <- habtools::circularity(shape)
```

```
  # dem metrics
  data$R_dem <- rg(dem, L0 = 1)

  data
}) %>% plyr::ldply()

# add growth form information from Zawada et al. (2019)
zawada <- readr::read_csv("data/3DLaserScannedColonies_MedToHighQuality_updatedFractalDimension.csv") %
  dplyr::rowwise() %>%
  dplyr::select(ScanID, GrowthFormVer, Species)


out$ScanID <- stringr::str_split_i(out$ScanID, pattern = fixed("PROW"), 1) %>%
  unlist() %>%
  str_split_i( "PROM",  1) %>%
  str_split_i( "PROF",  1) %>%
  str_split_i( "PRON",  1)

data <- inner_join(out, zawada)
```

In total, we quantified metrics for 84 colonies of various shapes. For simplicity, we subset and simplify growth form categories.

We can now visualize how metrics calculated from the meshes, DEMs, and 2D shapes compare.

```
ggplot(data) +
    geom_abline() +
    geom_smooth(aes(x = D3_dem, y = D3_mes),
                method = "lm", se = F, color = "black", linetype = 3) +
    geom_point(aes(x = D3_dem, y = D3_mes,
                   color = gf, shape = gf)) +
    scale_shape_manual(values = c(1:9)) +
    labs(x = "Fractal dimension (DEM)", y = "Fractal dimension (mesh)",
         color = "Growth form", shape = "Growth form") +

    ggplot(data) +
    geom_abline() +
    geom_smooth(aes(x = R_dem, y = R_mes),
                method = "lm", se = F, color = "black", linetype = 3) +
    geom_point(aes(x = R_dem, y = R_mes,
                   color = gf, shape = gf)) +
    scale_shape_manual(values = c(1:9)) +
    labs(x = "Rugosity (DEM)", y = "Rugosity (mesh)",
         color = "Growth form", shape = "Growth form") +
    scale_x_log10() +
    scale_y_log10() +

    ggplot(data) +
    geom_abline(slope = 1, intercept = 1) +
    geom_smooth(aes(x = D2_shp, y = D3_mes),
                method = "lm", se = F, color = "black", linetype = 3) +
    geom_point(aes(x = D2_shp, y = D3_mes,
                   color = gf, shape = gf)) +
    scale_shape_manual(values = c(1:9)) +
```

```
labs(x = "Fractal dimension (2D)", y = "Fractal dimension (mesh)",
     color = "Growth form", shape = "Growth form") +

ggplot(data) +
geom_abline() +
geom_smooth(aes(x = circ_shp, y = spher_mes),
            method = "lm", se = F, color = "black", linetype = 3) +
geom_point(aes(x = circ_shp, y = spher_mes,
               color = gf, shape = gf)) +
scale_shape_manual(values = c(1:9)) +
labs(x = "Circularity (2D)", y = "Sphericity (mesh)",
     color = "Growth form", shape = "Growth form") +

plot_layout(guides = "collect") + plot_annotation(tag_levels = "A") &

scale_color_fish_d("Callanthias_australis") &
theme_classic() + theme(legend.position = "right", text = element_text(size = 10))
```

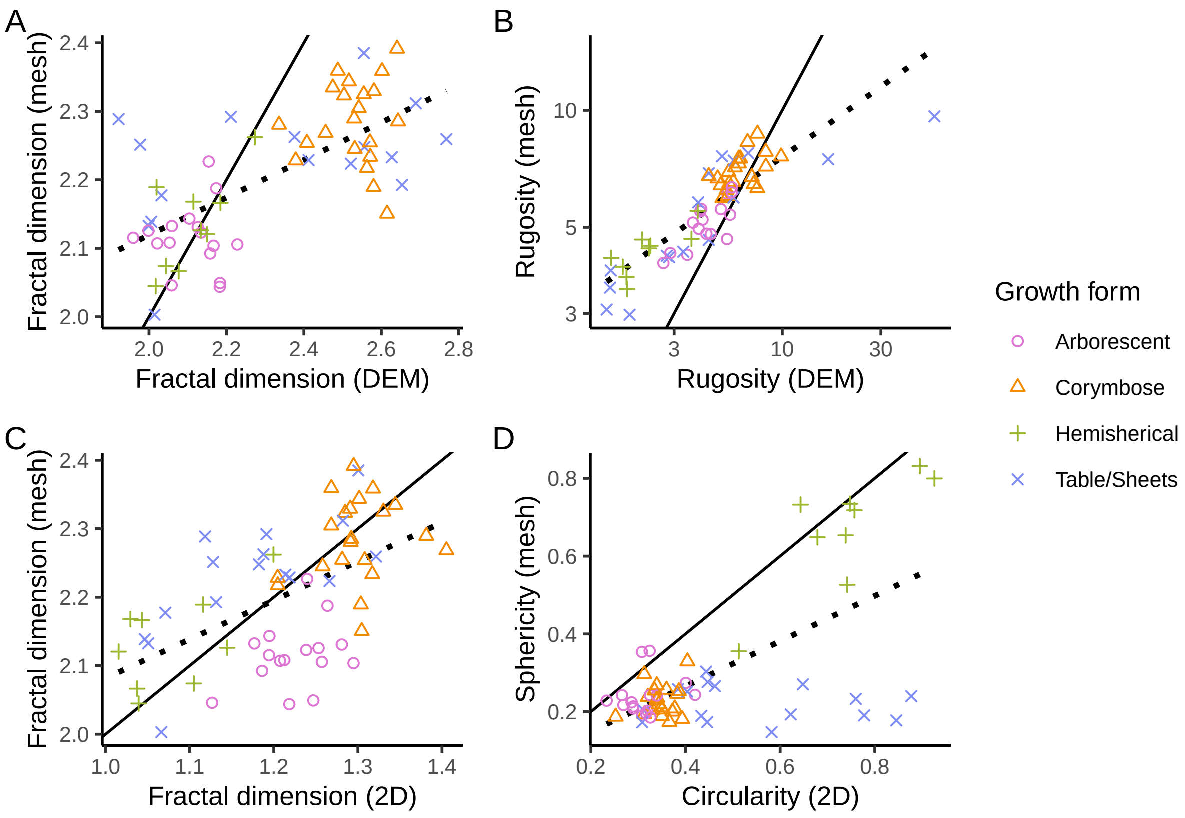
